# Evidence of dengue virus transmission and a diverse *Aedes* mosquito virome on the Democratic Republic of Congo-Angola border

**DOI:** 10.1101/2025.01.16.633031

**Authors:** Wenqiao He, Thierry Bobanga, Anne Piantadosi, Zachary R. Popkin-Hall, Fabien Vulu, Matthew H. Collins, Melchior M. Kashamuka, Antoinette K. Tshefu, Jonathan J. Juliano, Jonathan B. Parr

**Affiliations:** Institute for Global Health and Infectious Diseases and Division of Infectious Diseases, Department of Medicine, University of North Carolina at Chapel Hill, Chapel Hill, NC, United States; Department of Tropical Medicine, Faculty of Medicine, University of Kinshasa, Kinshasa, Democratic Republic of the Congo; Department of Pathology and Laboratory Medicine, Emory University School of Medicine, Atlanta, GA, USA; Division of Infectious Diseases, Department of Medicine, Emory University School of Medicine, Atlanta, GA, USA; Kinshasa School of Public Health, Kinshasa, Democratic Republic of the Congo

**Keywords:** *Aedes* mosquito, dengue, Democratic Republic of Congo, viral metagenomic

## Abstract

*Aedes* mosquitoes are widely distributed across the Democratic Republic of Congo (DRC), and are major vectors of dengue (DENV), Zika, chikungunya (CHIKV), and yellow fever (YFV) viruses. While the high burden of malaria in the DRC receives considerable attention, arboviruses remain understudied. In the setting of recent CHIKV and YFV outbreaks in southwestern DRC, we collected *Aedes* mosquitoes in three areas of Kimpese, DRC, near the Angola border, to investigate their virome. Metagenomic and targeted sequencing of eight randomly selected field mosquito pools, comprising 155 mosquitoes from three collection sites, confirmed high-confidence DENV reads and human blood meals in six (75%) and eight (100%) pools, respectively. We find diverse mosquito viromes including other known and putative human and animal viruses. Our findings provide strong evidence of endemic DENV transmission along the DRC-Angola border and illustrate the potential of wild-caught mosquitoes for xenosurveillance of emerging pathogens.

## Introduction

Arthropod-borne pathogens, including viruses, bacteria, and parasites, account for more than 17% of all infectious diseases ^1,2^, and over 700 arboviruses have been documented to infect humans across a wide geographic distribution ^3,4^. The number of outbreaks and endemic infections caused by emergent and re-emergent mosquito-borne arboviruses has increased worldwide over the last two decades ^5–7^. *Aedes* mosquitoes are particularly important vectors for multiple important human pathogens, including dengue virus (DENV), Zika virus (ZIKV), yellow fever virus (YFV), and chikungunya virus (CHIKV) ^8–11^. The warm climate and suitable environmental conditions in tropical regions support the proliferation of both mosquitoes and the viruses they transmit. As a result, tropical regions tend to have higher diversity and prevalence of arboviruses than other regions. However, arbovirus transmission is understudied across Africa, where improved understanding is urgently needed to guide the rollout of newly available DENV vaccines, in particular ^12^.

The Democratic Republic of the Congo (DRC) is the largest country in central Africa with a population of over 100 million, and a tropical climate suitable for mosquitoes. Approximately 248 mosquito species are thought to reside in the DRC ^13^, including 17 *Aedes* species, among which *Aedes aegypti* and *Aedes albopictus* are two major disease vectors ^7,14^. A recent survey found that *Ae. albopictus* is expanding its distribution in the DRC and displacing native *Aedes* species ^15^. Chikungunya, YFV, and DENV outbreaks and infections are known to occur in the DRC ^16–18^, and increasing reports of mosquito-borne and emerging viral diseases have been recorded ^19^. Considerable attention is devoted to the high burden of malaria ^20^ in the DRC, but arboviruses remain neglected, with limited studies of humans and mosquitoes to-date ^3,17–19,21^. The lack of diagnostic tools and surveillance systems for these diseases hinders our understanding of their impact and epidemiology ^22^.

Metagenomic sequencing approaches can be used for identification of viral DNA and/or RNA within diverse sample types. These methods have recently been applied to mosquito virome analyses, particularly in the context of mosquito-borne pathogen transmission and xenosurveillance, to enhance understanding of host–vector–environment interactions and support integrated monitoring of pathogens and exposures at the interface of animal and public health ^23–26^. To improve our understanding of the role of *Aedes* mosquito vectors in arboviral transmission in the DRC, we collected and sequenced the virome of mosquito pools from three areas of Kimpese, a health zone in Kongo Central province in southwestern DRC near the Angola border, a region that has experienced recent arboviral outbreaks ^3,16^ (**Figure 1**). We observed diverse viromes and identify strong evidence of active DENV transmission in the region, corroborated by viral sequences, DENV PCR, and mosquito blood meal analysis.

**Figure 1.**
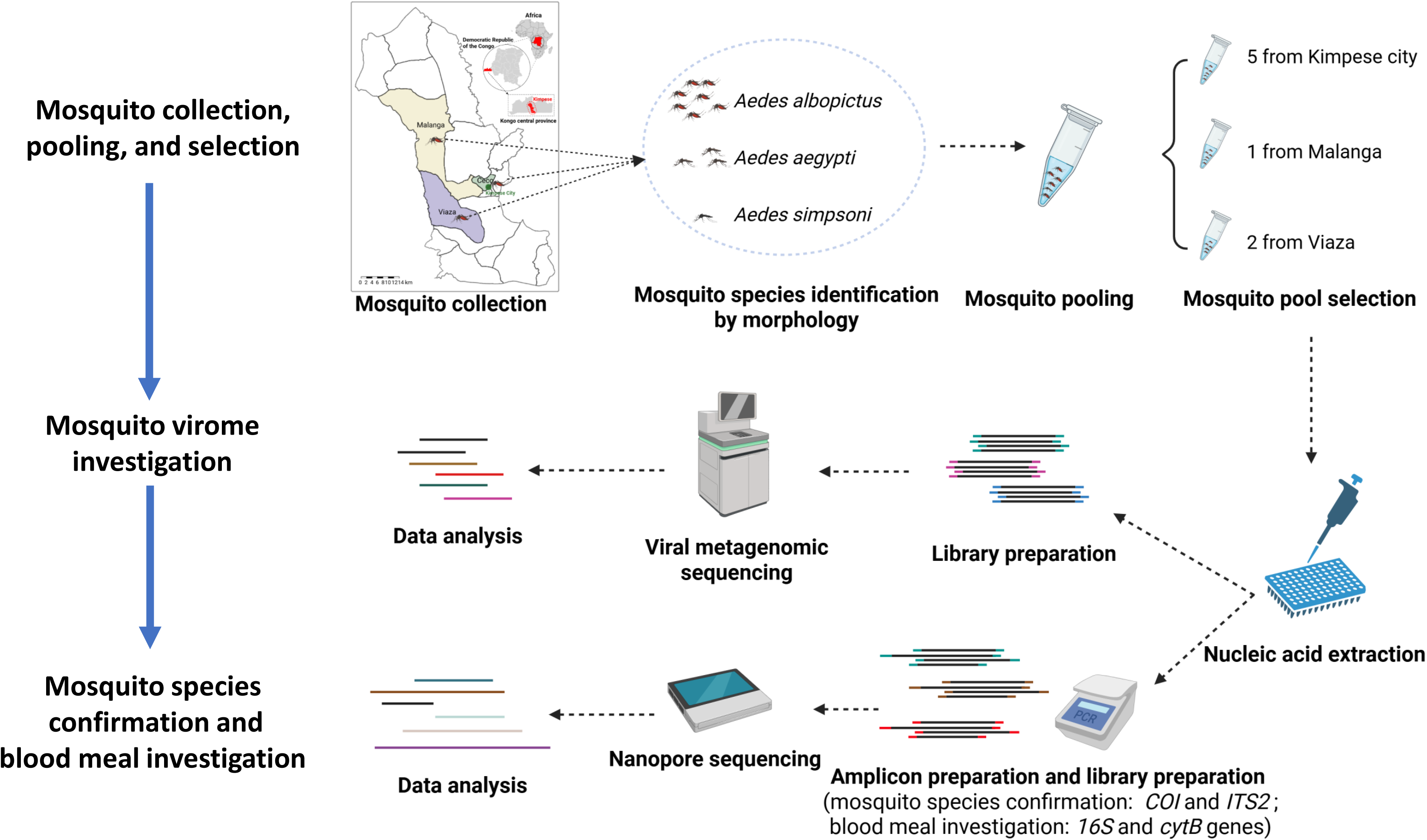
Schematic of the viral metagenomic, mosquito species, and blood meal analysis of *Aedes* mosquitoes from the Democratic Republic of the Congo. Created in BioRender.

## Results

### Mosquito collection and species confirmation

More than 600 adult *Aedes* mosquitoes **(Table S1)** were collected and distributed into 37 pools based on sample collection sites (21 pools from Kimpese city, 6 pools from Malanga, and 10 pools from Viaza, **Figure 2a**). Three species of *Aedes* mosquitoes were identified by morphology: *Aedes albopictus (n=653, 98.34%)*, *Aedes aegypti (n=8, 1.20%)*, and *Aedes simpsoni (n=3, 0.45%)*. Among the eight sequenced pools (**Figure 2b**), all contained a majority of *Ae. albopictus* (**Table S2**) based on their *ITS2* and *COI* gene sequences.

**Figure 2.**
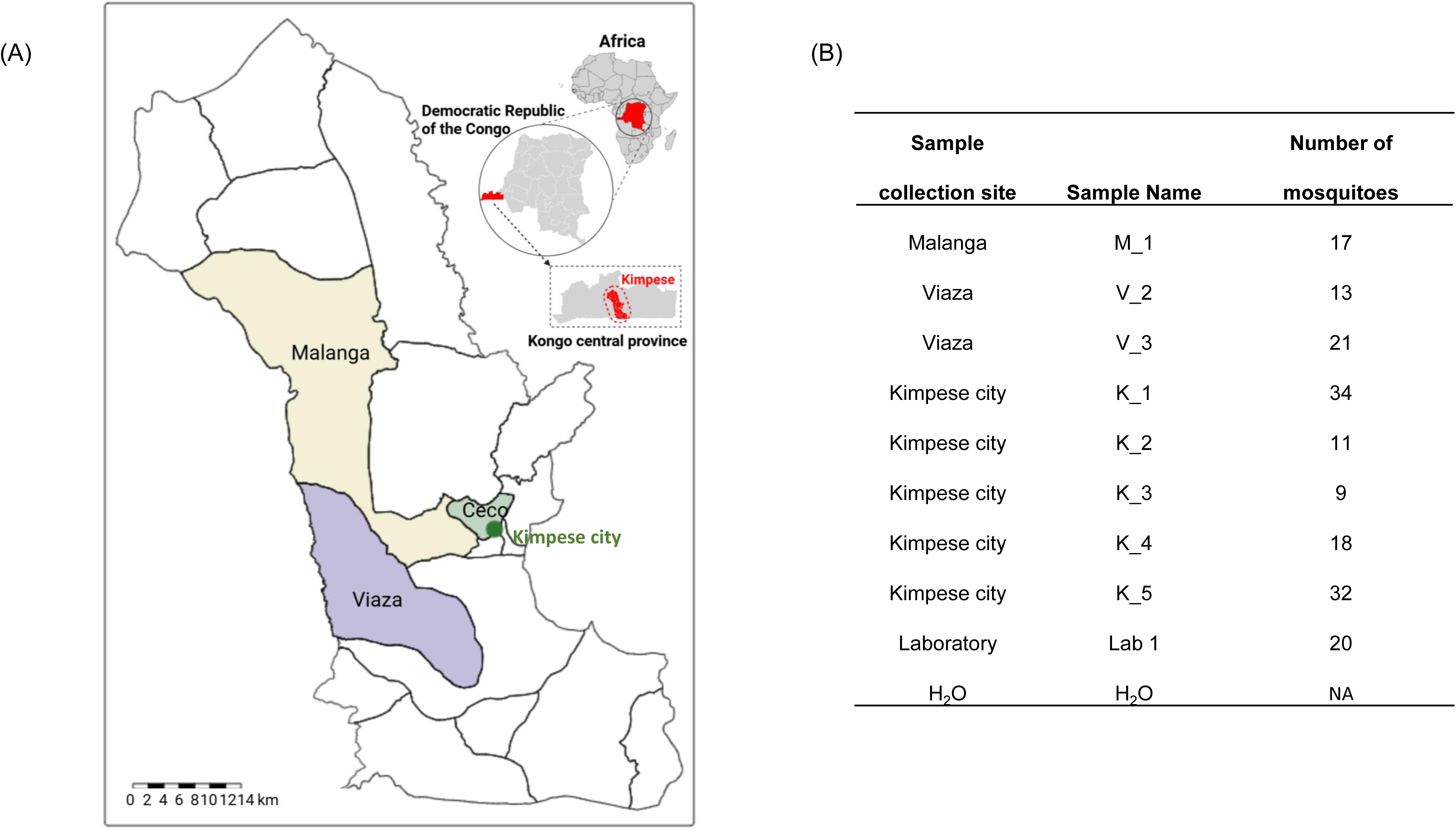
*Aedes* mosquitoes trapped in the DRC. A) Sample collection areas in Kimpese, DRC. Kimpese city is annotated by a green dot. B) Detailed information of the selected mosquito pools, including health areas where they were collected, and including laboratory-reared *Ae. aegypti* (Lab 1) and water (H_2_O) controls.

### Sequencing output

A total of 3,795,246,904 raw reads were obtained by Illumina sequencing. A range of 78.2-84.4% paired-end reads in each field mosquito pool were successfully merged. After filtering out low quality reads and those mapping to host and contaminating contigs (**Figure 3a and Table S3**), 185,621,738 reads remained, of which 144,469,836 were unclassified reads (66.2% of total) that did not map to any sequences in the KrakenUniq default nt database. Classified reads were mapped to known viruses, bacteria, eukaryota, and archaea.

**Figure 3.**
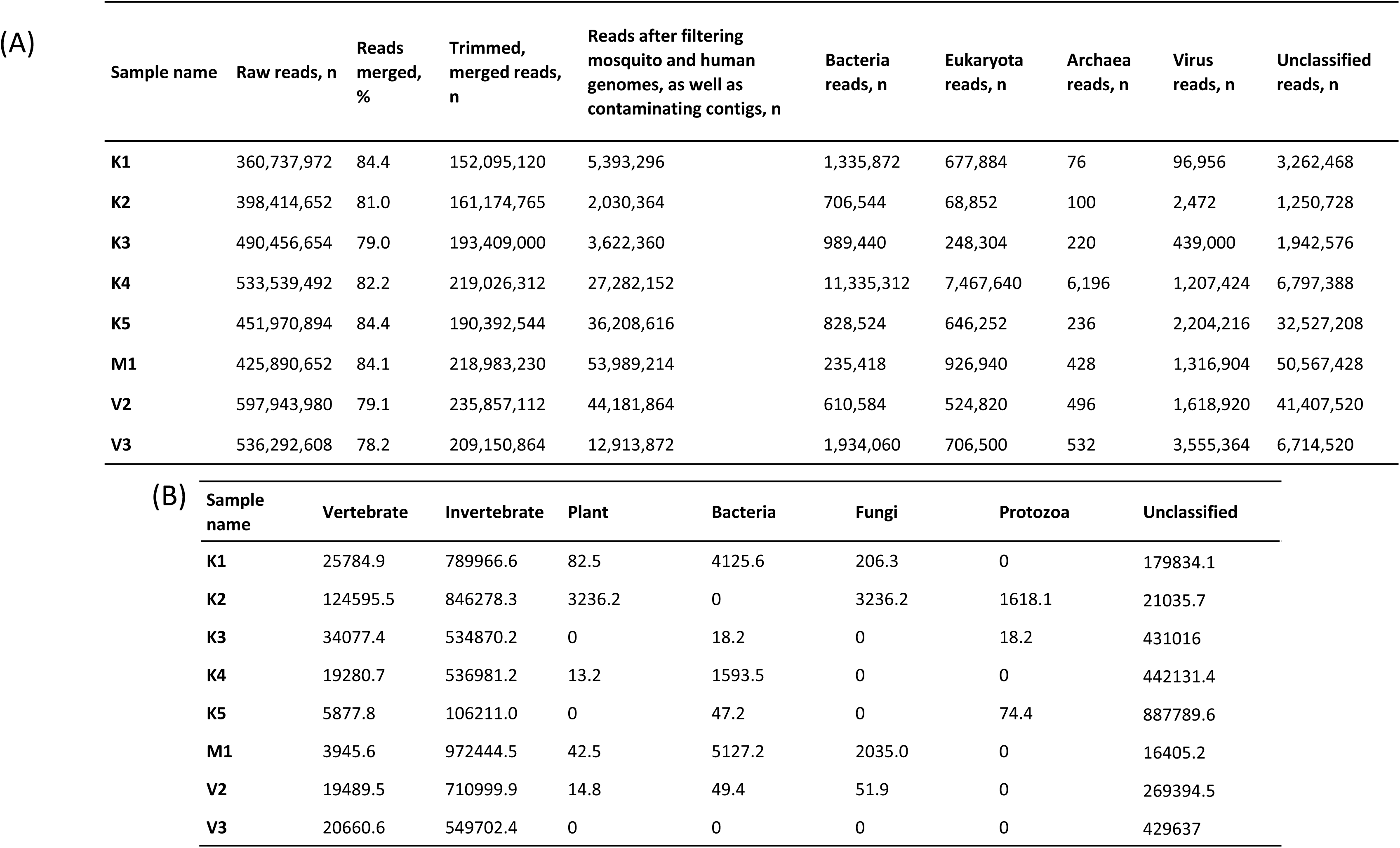
Summary of read counts based on the KrakenUniq nt database. **A)** Summary statistics for sequenced mosquito pools. **B)** Virus reads by presumed host, reported as reads per million based on KrakenUniq annotation using the nt database.

### Viral metagenomic analysis

Results confirm a highly diverse virome in wild-caught *Aedes* from the DRC, with viruses specific to invertebrates, vertebrates, plants, bacteria, fungi, and protozoa, as well as an unclassified group of viruses (**Figure 3b**). We do not observe a correlation between the number of viral taxa and the number of mosquitoes in the field-collected mosquito pools.

Most of the viral genera detected in the laboratory-uninfected *Ae. aegypti* mosquito pool and the water control are not found in the field mosquito pools (**Table S4**). Although reads annotated as viral genera *Betacoronavirus*, *Mimivirus*, and *Pahexavirus* are detected in both the water control and some field mosquito pools, BLAST results show no true matches to *Betacoronavirus* in the water control and reveal that the *Mimivirus* and *Pahexavirus* reads in field mosquito pools mapped to different genomic regions than those in the water control; therefore, no reads in field mosquito pools are removed in this filtering step. Based on taxonomic classification using the KrakenUniq with the NCBI-nt database, and after applying the RPM < 10 threshold and index-hopping filter, we identify 33 unique total viral families and 28 unique total viral genera in the field mosquito pools, with each pool containing between 3-17 viral families and 2-16 viral genera (**Figure 4, S1, and S2, Supplementary Material**).

**Figure 4.**
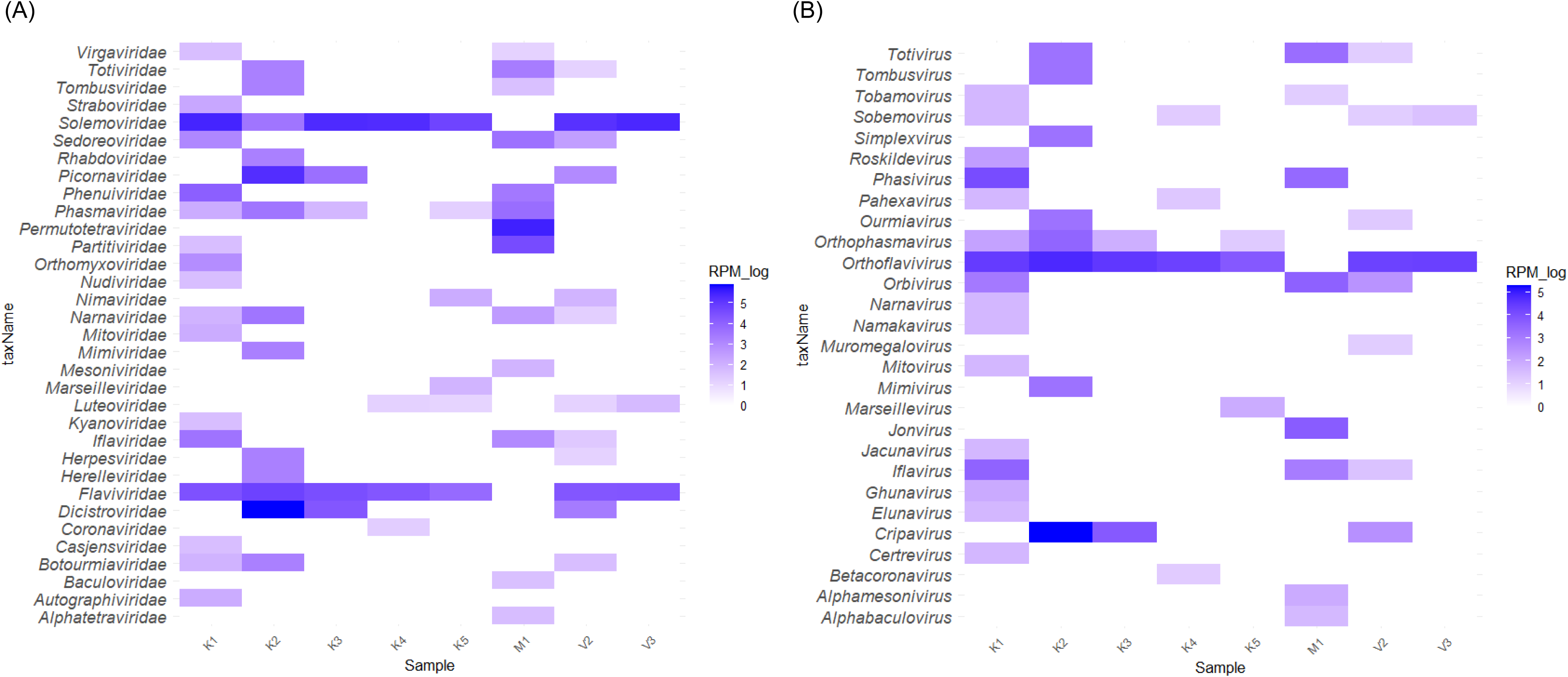
Viral families and genera in field mosquito pools. **A)** Heatmap based on reads per million viral reads in each mosquito pool (family level). **B)** Heatmap based on reads per million viral reads in each mosquito pool (genus level). Per International Committee on Taxonomy of Viruses recommendations, reads of *Orthoflavivirus* comprises both *Orthoflavivirus* and *Flavivirus* identified by KrakenUniq.

*Flaviviridae* is found in all field mosquito pools collected from Kimpese city and Viaza. *Orthoflavivirus,* which includes viruses formerly classified under the *Orthoflavivirus* and *Flavivirus* genera ^27^, is the most abundant viral genus in most Kimpese mosquito pools (except K2) and all Viaza pools, with a large number of reads mapping to *Orthoflavivirus denguei*. K2 contains a large number of reads annotated as genus *Cripavirus* (family *Dicistroviridae*). *Jonvirus*, a genus typically associated with insect viruses, shows the highest relative abundance in the mosquito pool from Malanga and is exclusively detected in that pool.

Several viral genera capable of infecting humans and animals are detected in mosquito pools collected from Kimpese city. These include *Betacoronavirus* (family *Coronaviridae*), *Marseillevirus* (family *Marseilleviridae*), and *Simplexvirus* (family *Herpesviridae*). Genus *Orthophasmavirus* is found exclusively in Kimpese pools, while *Alphabaculovirus* and *Alphamesonivirus* are detected only in the mosquito pool from Malanga. These genera contain insect-associated viruses. *Muromegalovirus*, a genus within the family *Herpesviridae* known to naturally infect rodents, is detected exclusively in one Viaza pool and is not in mosquito pools from Kimpese city and Malanga. All corresponding reads map to viral species *Muromegalovirus muridbeta2*.

PCA based on detected viral genera to explore clustering by sample collection area is shown in **Figure S3 and Table S5**. Geographical clustering is observed among samples from Kimpese city and Viaza. Notably, two mosquito pools from Kimpese city differ substantially from the other three pools collected at the same site. Small sample sizes at each sample collection site require this PCA to be interpreted as an exploratory visualization with insufficient precision to identify statistically robust differences within or between sites.

KrakenUniq annotation results prior to applying the RPM < 10 threshold and index-hopping filter are provided in the Supplementary Notes. These datasets include all viral sequences remaining after initial quality filtering steps, such as removal of low-quality reads, host-derived sequences, and contaminant contigs or reads, providing a comprehensive overview of the detected viral diversity. Notably, some pathogens and potential pathogens were detected with low read counts, including *human pegivirus* and *human blood-associated dicistrovirus*.

### Analysis of reads mapping to pathogens and potential pathogens

BLAST-confirmed reads mapping to viral species related to human or animal diseases are detected in multiple mosquito pools, including DENV and *bat faecal associated dicistrovirus 4* (**Figure 5**). Though SPAdes does not successfully assemble any contigs spanning multiple DENV reads in this study, BLAST-confirmed, merged paired-end reads mapping to DENV are found in 6 out of the 8 (75%) field mosquito pools (**Table S6**). Consensus sequences generated from the BLAST-confirmed reads mapping to DENV-2 in one mosquito pool (K5) is highly similar to one sequence from Malaysia (MH048672.1 10664-10770, 107 bp, nucleotide identity = 94%, **Table S6**). Consensus sequences mapping to DENV-4 found in all six positive pools are short (< 100bp) and show high similarity to the 5’-untranslated (UTR) region of a published sequence from Thailand (MG601754.1, nucleotide identity ≥ 97.78%) (**Table S6**). Amplification of partial envelope protein E gene, RNA-dependent RNA polymerase NS5 gene, and fragments adjacent to the metagenomic consensus sequences using published assays failed ^28–30^, likely due to low DENV RNA concentration and RNA degradation within mosquito pools (median peak RNA fragment length 176 nt, IQR 115-178). We note that these DENV-4 reads also map to *Wenzhou sobemo-like virus 4* sequences, but with lower similarity (nucleotide identity ≤ 96.67%) and higher e-values. We find high similarity between consensus sequences mapping to *bat fecal-associated dicistrovirus 4* to a published *bat faecal associated dicistrovirus 4* sequence detected in feces from *Pteropus poliocephalus* from Australia (ON872534.1, 7078-7330, 253 bp) in three mosquito pools (**Figure 5**). However, we are unable to assemble contigs for this virus, and resolution of phylogenetic analysis is therefore limited. Additional details on these viral reads are provided in **Figure 6** and **Table S7**.

**Figure 5.**
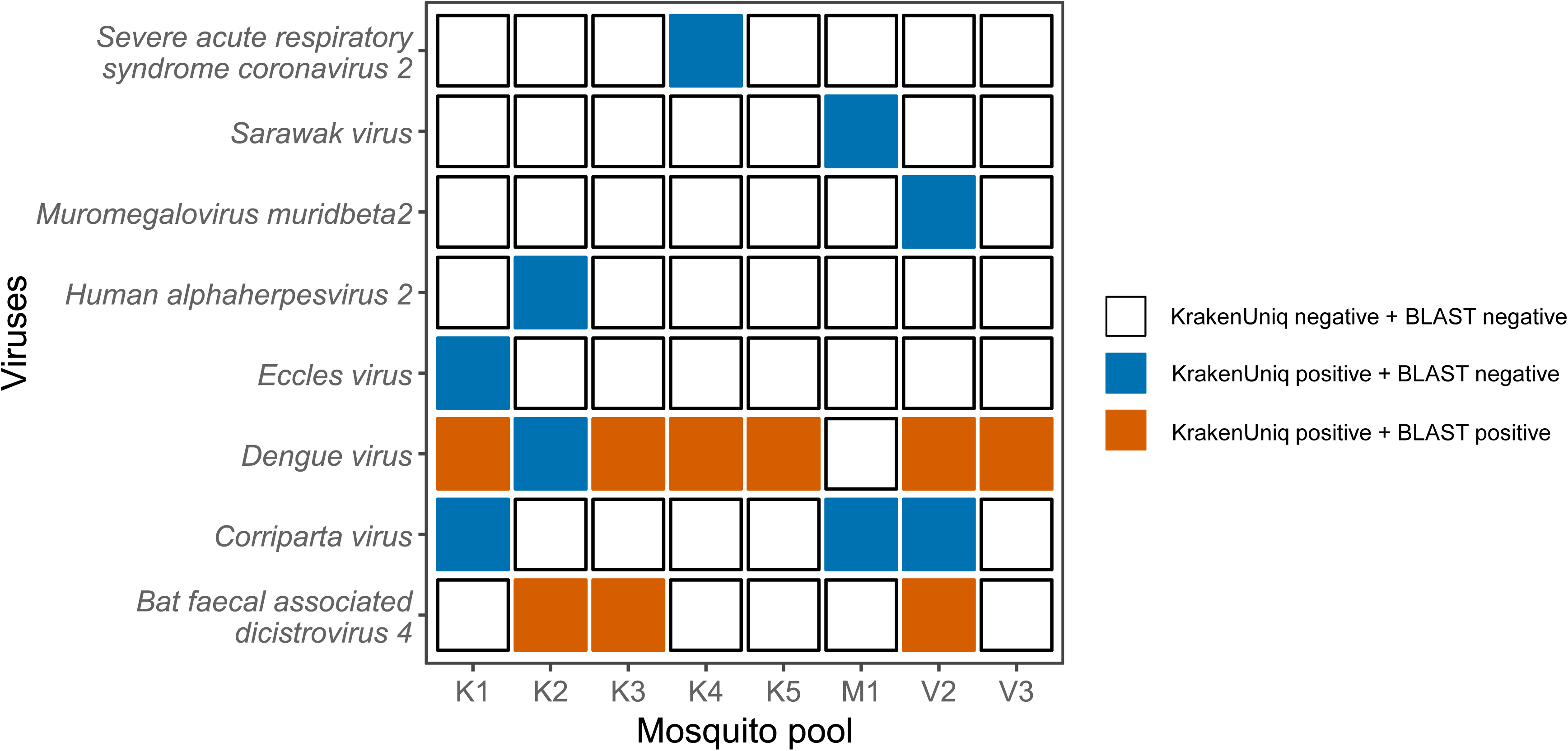
Known human or animal viruses detected in mosquito pools. Annotation results based on KrakenUniq with default nt database and BLAST are shown in different colors. Vermillion rectangulars signify confirmation by two methods used.

**Figure 6.**
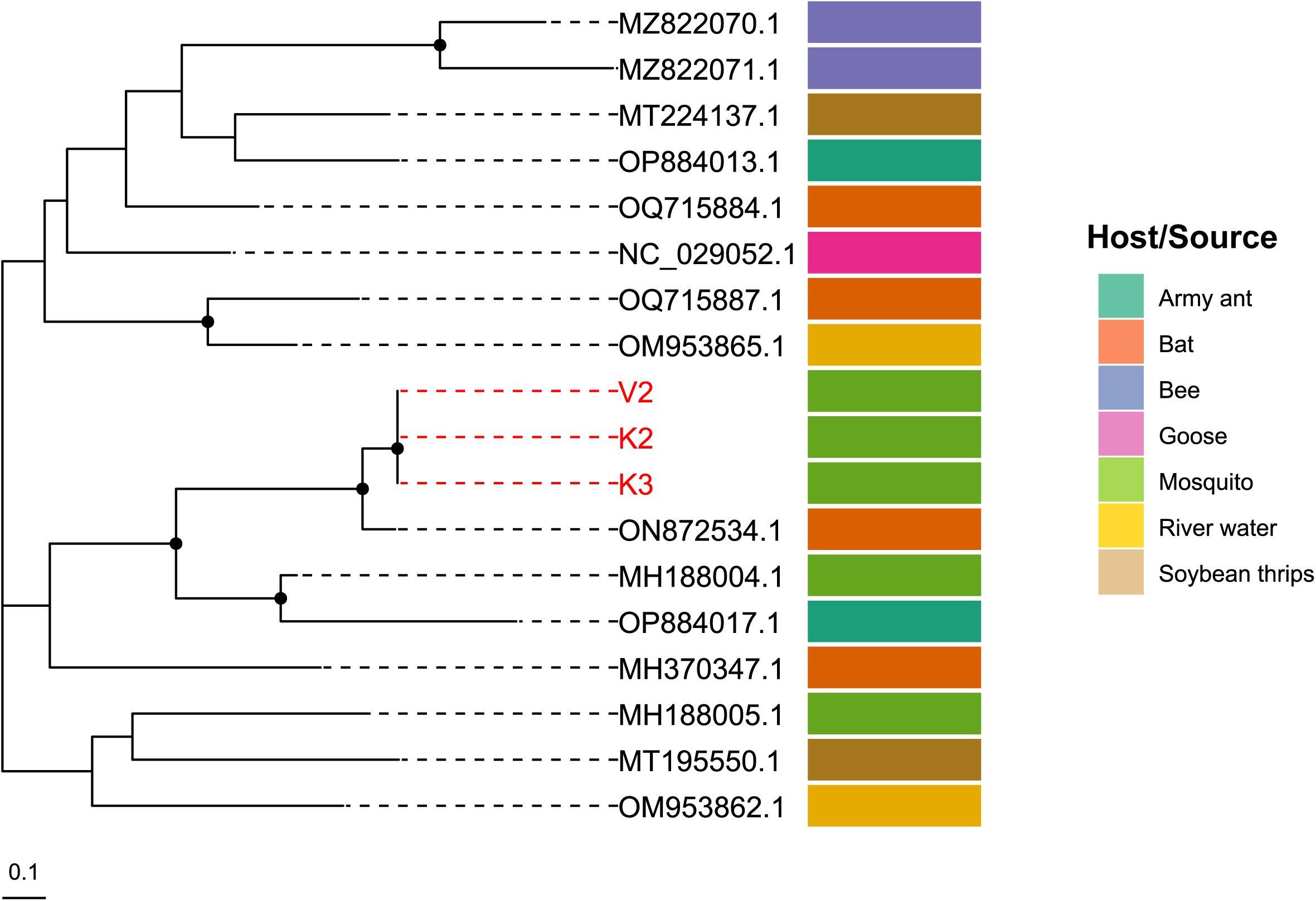
**Phylogenetic analysis of bat faecal associated dicistrovirus 4** reads in DRC field mosquito pools (red) mapping to the non-coding region (7078 - 7330 nt based on ON872534.1) compared to published sequences (black), constructed using RAxML-NG with the HKY+F+G4 model. Black dots indicate nodes with bootstrap support greater than 80% across 1,000 replicates.

### PCR confirmation of DENV and mosquito blood meal investigation

Real-time PCR targeting the DENV polyprotein gene, distinct from genomic regions identified in our metagenomic analysis, confirms DENV RNA in the three mosquito pools with highest DENV read counts in metagenomic analysis (**Table S6**), indicating consistency between next-generation sequencing and real-time PCR within its limits of detection.

Investigation of mosquito blood meals using amplicon sequencing showed that mosquitoes from all selected pools had taken blood from *Homo sapiens* (**Figure 7**). Additionally, we detected reads mapping to non-human primates (4/8 pools), rodents (3/8 pools), canines (3/8 pools), and sheep (1/8 pools).

**Figure 7.**
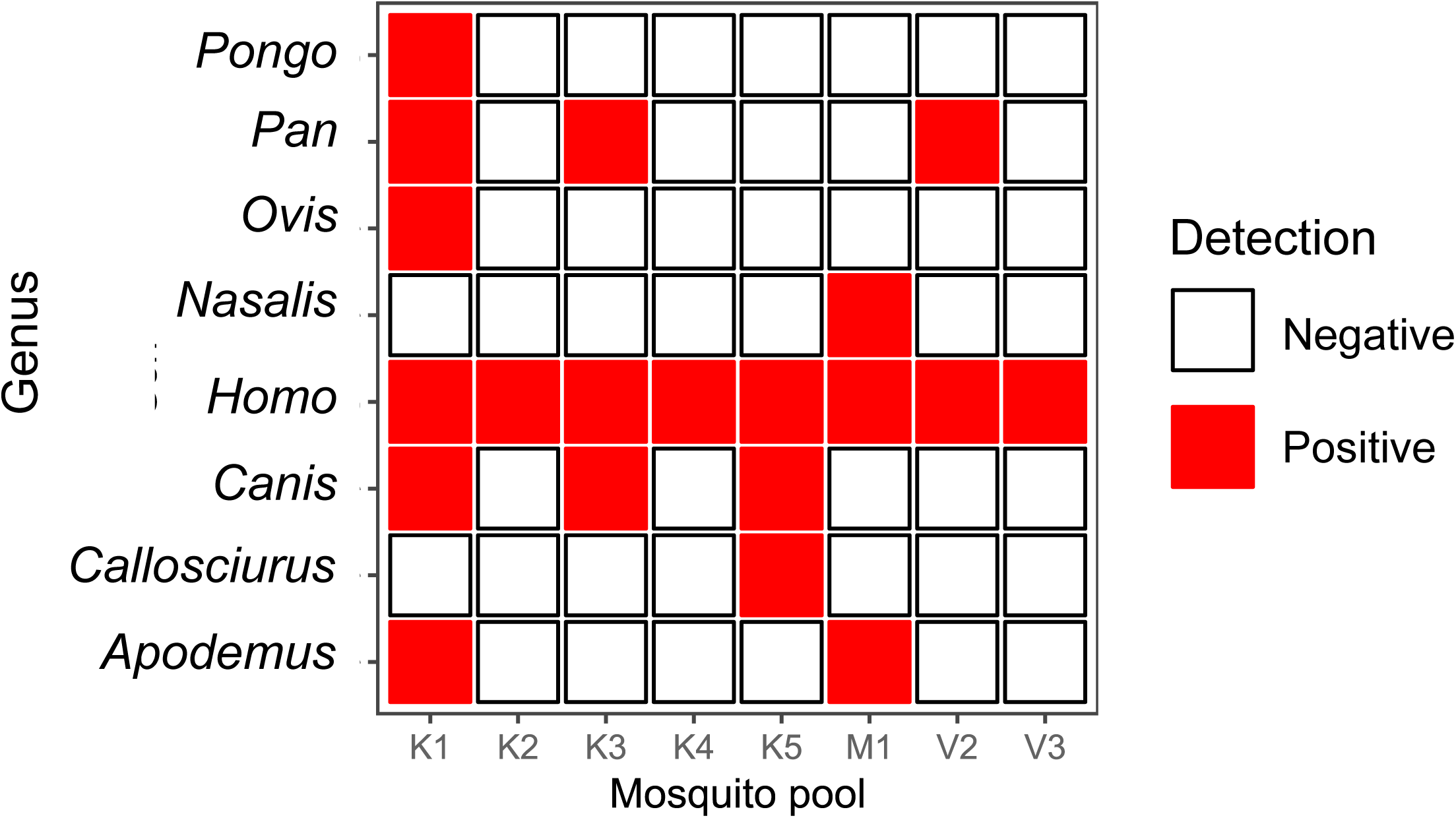
**Mosquito blood meal sources** determined by amplicon sequencing of the 16S ribosomal RNA and cytochrome b genes.

### Insect-specific viruses

We find a large number of insect-specific viruses in field mosquito pools, with most belonging to the families *Solemoviridae* and *Flaviviridae*. *Wenzhou sobemo-like virus 4*, *Hubei mosquito virus 2*, *Guangzhou sobemo-like virus*, *Sichuan mosquito sobemo-like virus*, and *Aedes flavivirus* are the most prevalent species (**Table S8**). Except for *Wenzhou sobemo-like virus 4* (6/8, 75%), the other four insect-specific viruses are detected in 7 out of 8 (87.5%) mosquito pools with high relative abundance.

BLAST-confirmed reads of *Aedes flavivirus* were detected in 7 mosquito pools, whereas reads of *Wenzhou sobemo-like virus 4*, *Guangzhou sobemo-like virus*, and *Sichuan mosquito sobemo-like virus* were detected in 6 pools. Consensus sequences were successfully generated based on the BLAST confirmed reads of these four insect-specific viruses. A large number of reads initially classified as *Hubei mosquito virus 2* by KrakenUniq were reassigned to *Guangzhou sobemo-like virus* after BLAST confirmation, likely attributable to the high sensitivity but lower specificity of the metagenomic assay. Only one mosquito pool (M1) has BLAST-confirmed *Hubei mosquito virus 2* reads. Though the consensus sequences were short (< 150 bp), they showed high similarity (≥ 94.19%) to multiple regions to a published *Hubei mosquito virus 2* sequence from China (MW452307.1, 72-136, 64bp; 140-226, 86bp; 227-318, 91bp; and 3390-3519, 129bp) using BLAST.

Phylogenetic analysis based on the consensus sequences shows that *Aedes flavivirus Wenzhou sobemo-like virus 4*, *Guangzhou sobemo-like virus*, and *Sichuan mosquito sobemo-like virus* detected in *Aedes* mosquitoes from DRC are highly similar to the corresponding published sequences from China (**Figure S4-S7**).

## Discussion

Our findings confirm the presence of dengue and other viruses with human or animal pathogenic potential in wild-caught *Aedes* mosquitoes circulating near the DRC’s border with Angola. The presence of human blood in all mosquito pools strongly supports the possibility that there is active dengue transmission in the Kimpese region, including peri-urban and rural zones outside of the city. The combination of a diverse mosquito virome and the presence of blood meals from multiple mammals suggests robust opportunities for arthropod-borne viral transmission between species, as well as for spillover events. Together, our experience suggests a future role for xenosurveillance and metagenomic sequencing as a complement to existing data systems developed to monitor for emerging infectious diseases in countries like the DRC.

Multiple arboviral disease outbreaks have been documented in the DRC during the last century, including yellow fever, dengue, chikungunya, and others ^3,17,18,21^. Except for DENV, we did not find other known human pathogenic arboviruses in the selected *Aedes* pools. In practice, mosquito xenosurveillance approaches should be tailored to specific use cases. Given the typically low detection rates of many arboviruses in mosquito populations, longitudinal sampling across multiple transmission seasons, increased spatial coverage within ecologically diverse sites can be used to improve detection of known and emerging pathogens. Repeated sampling over time, combined with targeted mosquito sampling, pathogen-specific PCR or sequencing, during periods of increased vector abundance or reported human or animal cases, is likely to substantially increase detection sensitivity. In this context, metagenomic sequencing is best suited as a complementary tool within an integrated surveillance strategy.

Detection of DENV in mosquito pools by RT-PCR suggests active DENV circulation in the Kimpese region, and metagenomic sequencing reads from small regions of the genome point towards the circulation of both DENV-2 and DENV-4. Three of the four DENV serotypes ^31^ have been recorded in the human population of the DRC (DENV-1, DENV-2, and DENV-3), with DENV-1 being the most frequently reported ^3^. One study in Singapore reported that the majority of the dengue hemorrhagic fever cases involving DENV-2 were from the cosmopolitan genotype ^32^. DENV-2 detected in our study is most similar to the cosmopolitan genotype of DENV-2, which has been previously reported in the DRC ^33^. Although the DENV-4 reads detected in our study also map to *Wenzhou sobemo-like virus 4*, an insect-specific virus, they have lower similarity (nucleotide identity ≤96.67%) and higher e-values. Our study provides preliminary evidence that DENV-4 may circulate in *Aedes* mosquitoes from the DRC and therefore infect people in this region.

Data from all *Aedes* mosquito pools in our study confirm human blood meals. Previous studies have reported high dietary diversity of *Aedes* mosquitoes, with a predominant preference for mammalian hosts ^34,35^. We corroborate these findings, with diverse mammalian blood meals observed across mosquito pools, including blood from non-human primates. Non-human primates can be amplifying hosts and play considerable roles in the sylvatic cycle of DENV transmission, which can spark DENV epidemics in nearby human populations ^36^. Detection of rodent, canine, and sheep genomes in the collected mosquito pools illustrate the diversity of hosts of *Aedes*, though further research is needed to confirm the roles of these hosts in DENV transmission, including the contribution to human infection. Notably, three (K2, K4, and V3) mosquito pools appear to have exclusively fed on human blood.

Similar windows of detection have been observed for pathogen RNA and human DNA in mosquitoes following a blood meal. Past work has shown complete digestion of a human blood meal varies depending on the mosquito species and typically takes about 36 hours to 3 days ^37,38^. The persistence of pathogen RNA or DNA in mosquitoes can vary depending on the pathogen type, viral strain, viral load, and mosquito species. Non-arboviral RNA can remain detectable in the mosquito midgut for a field-relevant time-frame—up to 18 hours for west nile virus RNA and up to 24 hours for HIV-1 RNA after blood feeding ^39^. Influenza virus RNA is detectable up to 36 hours post feeding ^38^. For arboviruses, the detection window may be longer. For example, one study showed a reduction in DENV-2 RNA signal as early as 12 hours post-feeding, with little to no signal detected after 24 hours. However, if the virus successfully replicated in mosquito tissues, a subsequent increase in RNA levels was observed between 100 and 200 hours post-blood meal ^38^. The co-detection of human DNA and DENV RNA in the same mosquito pools suggests a common source, likely a viremic human host. Although detection of DENV RNA does not necessarily indicate the presence of infectious virions, its presence in two of these pools (K4 and V3) strongly suggests active DENV circulation in these communities.

In addition to DENV, we identify reads mapping to *bat faecal associated dicistrovirus 4*. Detection of this non-arbovirus might be related to incomplete degradation of the viral material by the mosquito digestive system at the time of sampling. This is the first report of *bat faecal associated dicistrovirus 4* in the DRC. While dicistrovirus is believed to exclusively infect invertebrates, it has been recently reported in blood samples from febrile patients in Tanzania and Nigeria ^40,41^, the intestinal content of a captive red squirrel with enteritis ^42^, and has been identified in free-living gorillas in neighboring Republic of Congo ^43^. Interestingly, we found *bat faecal associated dicistrovirus 4* reads in a mosquito pool (K2) that exclusively fed on human blood, suggesting that it came from an infected human. However, the host range and pathogenicity of these viruses are unclear.

Similar to other studies ^44,45^, insect-specific viruses were frequently detected in *Aedes* mosquitoes collected from the DRC, including *Aedes flavivirus*, *Wenzhou sobemo-like virus 4*, *Guangzhou sobemo-like virus*, and *Sichuan mosquito sobemo-like virus*. Although their implications for public health remain unclear, the high detection rates observed in *Aedes* mosquitoes highlight the need for further investigation into their ecological roles and potential interactions with pathogenic arboviruses. The sequences obtained for these viruses were highly similar to those previously reported from China. One possible explanation is widespread circulation of globally distributed viral lineages that may have dispersed between Asia and Africa through infected mosquitoes or eggs. This close genetic relatedness also suggests the possibility of long-term maintenance via vertical transmission within *Aedes* populations, with minimal local diversification despite substantial geographic separation.

Our findings also shed light on circulating *Aedes* species along the DRC-Angola border. Kimpese comprises a small city and surrounding region where more than 100,000 people are thought to live ^46^, though the DRC has not performed a formal census since the 1950s. It is located in the Kongo Central province, has a tropical climate, and spans mountains, valleys, and grasslands. *Aedes aegypti* and *Ae. albopictus* are the main vectors of several major pathogenic arboviruses, with *Ae. aegypti* being more competent for DENV and ZIKV transmission ^47^. *Ae. aegypti* was previously the dominant *Aedes* species in the DRC, until recent, rapid replacement of native *Aedes* species by *Ae. albopictus* in the western and northern regions of DRC ^15^. Our findings corroborate these changes in the DRC *Aedes* population; most collected *Aedes sp.* mosquitoes are *Ae. albopictus* (98.3%). The highly diverse viromes in our *Aedes* mosquito pools include viruses specific to invertebrates, vertebrates, plants, bacteria, fungi, and protozoa. These reads likely reflect the mosquitoes’ ingestion of blood, nectar, and environment substrates ^48^. However, the large number of unclassified reads within all pools confirms how much remains to be learned about this important mosquito population.

Kimpese, Viaza, and Malanga are all situated within Kongo Central Province but differ modestly in elevation (approximately 300m above sea level) and landscape context. Kimpese lies at a relatively higher and more dissected terrain, Malanga occupies slightly lower elevations, and Viaza is positioned within a comparable low–to–mid elevation zone. All three localities are characterized by predominantly rural–agricultural settings; however, Kimpese serves as a more established market and transport hub, whereas Viaza and Malanga represent smaller, village-based agricultural communities within a similar savanna–farmland mosaic. Although not statistically significant, the clustering observed among the samples from Kimpese city and Viaza in PCA indicates possible geographical structuring within local *Aedes* mosquito populations. Most of the top viral genera contributing to the variance in principal components 1 and 2 have been detected in insects, plants, and bacteria. These findings suggest that local ecological and environmental factors—such as elevation, land use patterns, or vegetation—may be shaping the mosquito virome. In addition, the variability observed between mosquito pools within Kimpese city may reflect microenvironmental differences or heterogeneous exposure to viral sources within an urban landscape.

Several limitations of our study should be highlighted. First, single-timepoint sampling of mosquito pools may have limited our ability to detect rare or sporadically circulating human- or animal-pathogenic arboviruses. Second, mosquitoes were not pooled based on morphological mosquito species, and our metagenomic analysis focuses on *Aedes*. Analysis of other mosquito genera would provide broader insights into differences in arboviral transmission and viromes in the DRC. Third, downsampling was necessary to accommodate sequencing costs. We sequence a small number of mosquito pools from sites within one region, which provides useful insights into arboviral transmission in Kimpese but limits the generalizability of our findings to other regions. Fourth, despite positive RT-PCR results and the identification of reads mapping to limited regions of the DENV genome, our attempts to amplify additional and longer DENV fragments using several published assays were unsuccessful, likely due to low RNA concentration and RNA degradation within the mosquito. As a result, we did not have sufficient genomic coverage to construct whole genomes or perform rigorous phylogenetic analysis. Nor can metagenomic sequencing tell us whether infectious virions are present or definitively determine dengue types in circulation. Fifth, the use of mosquito pools improves the yield of our sequencing efforts but prevents us from estimating the prevalence of viruses in individual mosquitoes. Finally, viral metagenomic techniques have high sensitivity but low specificity. To overcome this shortcoming, we employ a conservative approach leveraging multiple bioinformatic tools complemented by real-time PCR, both of which confirm the presence of DENV in our sample set.

In conclusion, our study provides new insights into viruses infecting *Aedes* in the DRC and demonstrates the potential of wild-caught mosquito xenosurveillance. We use virome and blood-meal analysis to improve understanding of *Aedes* mosquito feeding behavior in the DRC, and present evidence of endemic DENV transmission in the Kimpese region near the DRC’s border with Angola DRC. These findings are particularly relevant to ongoing discussions about where to offer newly approved dengue vaccines and indicate a need for larger human studies to define dengue epidemiology in the DRC and across Africa ^49^. They also suggest a role for sustained vector control efforts targeting *Aedes* in the DRC, extending beyond outbreak response, and highlight opportunities for future arthropod-borne disease studies in the region.

## Methods

### Mosquito collection

Adult *Aedes* mosquitoes were collected from three health areas in Kimpese, DRC, Ceco (Kimpese city), Malanga, and Viaza (**Figure 2a**), using electric aspirators (Prokopack Aspirator, John W. Hock, Gainesville, USA) and BG sentinel traps (Biogents Inc, Regensburg, Germany) baited with dry ice. Mosquitoes were collected from April 23 - 30, 2022, during the rainy season when mosquito density is high. All collections were performed outdoors, and GPS coordinates along with a brief description of collection sites were recorded.

Collected mosquitoes were morphologically identified to the species level following the Huang morphological key ^50^. *Aedes* species were pooled by sample collection site. All field mosquito pools were preserved in DNA/RNA Shield (ZYMO research, CA, USA) and stored at -80°C.

Uninfected *Aedes aegypti* mosquitoes (NR-48920) were obtained from the NIH/NIAID Filariasis Research Reagent Resource Center through BEI Resources. A random subset of 20 mosquitoes was pooled to serve as the laboratory uninfected *Ae. aegypti* control.

### Nucleic acid extraction

Eight field *Aedes* mosquito pools, comprising a total of 155 mosquitoes, were randomly selected from the three health areas: five from Kimpese city, one from Malanga, and two from Viaza (**Figure 2b**). These pools represent approximately 20% of the total mosquito pools from each region. The selected mosquito pools were washed with 70% ethanol and rinsed twice with molecular-grade water before removal of mosquito heads ^51^. Total RNA was extracted from mosquito bodies from the selected field mosquito pools, the laboratory un-infected *Ae. aegypti* mosquito pool from BEI Resources, and one water control using the Quick-RNA Tissue/Insect kit (ZYMO research, CA, USA) following the manufacturer’s instructions. DNase I treatment using DNase I Set (ZYMO research, CA, USA) was carried out during the RNA extraction process.

### Mosquito species confirmation and blood meal investigation

To confirm the mosquito species, two regions within the cytochrome c oxidase I (*COI*), and one region within the second internal transcribed spacer (*ITS2*) genes were amplified from RNA extracted from the mosquito pools using published primers and conditions as shown in **Table S9** ^52–54^. DNA-based methods have been widely used in mosquito blood meal analysis and provide reliable results ^55^. The 16S ribosomal RNA (*16S*) and cytochrome *b* (*cytB*) genes are commonly used due to their species-specific resolution ^56,57^. In this study, we amplified *16S* and *cytB* from RNA extracted from field-collected mosquito pools using published primers (**Table S9**) ^58,59^. All amplicons from the same mosquito pool were combined in equimolar amounts. Then, 68 ng of DNA from each pooled amplicon was used for library construction. Amplicon libraries were constructed using the Native Barcoding Kit 96 V14 and sequenced on a Flongle Flow Cell (R10.4.1) using the MinION platform (Oxford Nanopore Technologies, Oxford, United Kingdom).

### Ribosomal RNA depletion, library preparation, and viral metagenomic sequencing

To increase viral metagenomic sequencing efficiency, mosquito ribosomal RNA (rRNA) depletion was carried out before library preparation using the NEBNext RNA Depletion Core Reagent set (New England BioLabs, MA, USA) with published probes targeting *Aedes* mosquitoes ^60^. RNA fragment lengths after rRNA depletion were assessed for five pools using Agilent TapeStation 4150 with High Sensitivity RNA ScreenTape (Agilent, Waldbronn, Germany). Sequencing libraries were prepared and barcoded using the NEBNext Ultra II Directional RNA Library Prep Kit and the Multiplex Oligos for Illumina (96 Unique Dual Index Primer Pairs) kit (New England BioLabs, Massachusetts, USA), respectively. The barcoded libraries were sequenced using the NovaSeq S4 platform (Illumina, California, USA) at the High-Throughput Sequencing Facility (HTSF) at the University of North Carolina at Chapel Hill to generate 150 bp paired-end reads. The expected total yield was 4-5 billion reads.

### Dengue virus confirmation by real-time PCR

To confirm DENV calls made by sequencing, we tested RNA from all mosquito pools using a published real-time PCR assay to detect pan-DENV RNA with the SuperScript III Platinum One-Step qRT-PCR kit (Life Technologies, Carlsbad, CA, USA) ^61^.

### Data analysis

To investigate viruses in the mosquito pools, Illumina NovaSeq sequencing reads were demultiplexed into individual samples based on their barcode sequences. Paired-end reads were then merged into single reads based on overlap detection using `BBMerge` (version 38.96) ^62^, followed by adapter removal using `Trimmomatic` (version 0.36) with default parameters, and filtering short reads by setting the minimum length as 36 ^63^. Reads mapping to human (GRCh38.p14), *Ae. albopictus* (Aalbo_primary.1), and *Ae. aegypti* (AaegL5.0) genomes, as well as the *Ae. simpsoni* CO1 gene (KT881399.1) were removed using `bwa-mem2` (version 2.2.1) ^64^ and `SAMtools` (version 1.21) ^65^.

Reads remaining in the laboratory un-infected *Ae. aegypti* mosquito pool and water control pool were assembled into contigs using `metaSPAdes` (version 4.0.0) with default parameters ^66^. These contigs were considered as contaminating contigs. Reads in field mosquito pools were aligned to the contaminating contigs using `bwa-mem2` (version 2.2.1), and mapping reads were removed using `SAMtools` (version 1.21). Taxonomic classification of the remaining reads were performed using `KrakenUniq` (version 1.0.4) in default mode, with the standard NCBI-nt database downloaded on July 31, 2023 ^67^. Then, reads from field mosquitoes mapping to sequences annotated as viral genera present in the laboratory uninfected *Ae. aegypti* mosquito pool and water control were excluded from the analysis. A viral genus or family was considered present by KrakenUniq if it had at least 10 mapped reads per million viral reads based on the KrakenUniq reports using the default NCBI-nt database ^26^. If the total read count of a specific viral genus or family in a specific library is less than 0.1% of the highest read count for that viral genus or family within the same sequencing lane, then it was considered as a false-positive due to index-hopping ^68^. The relative abundance of viral genera and families was calculated based on the KrakenUniq reports with default NCBI-nt database. This was done by dividing the number of reads mapping to specific taxonomic groups by the total number of viral reads, then multiplying the result by 100. Reads classified as viruses were extracted for further analysis. Principal component analysis (PCA) was performed based on the viral genus annotation.

BLAST (version 2.14.1), an alignment-based method, was used to confirm the classification of reads mapping to human and animal pathogens, as well as potential pathogens. BLAST-confirmed reads mapping to human and animal pathogens or potential pathogens were assembled into contigs using `metaSPAdes` (version 4.0.0) with default parameters. Consensus sequences were generated from merged paired-end reads aligned to the closest known relative reference sequence identified by BLAST. For phylogenetic analyses, consensus sequences were aligned with representative reference sequences (**Table S10-S16**) from NCBI Virus, RefSeq, as well as GenBank databases, including diverse genotypes/serotypes spanning multiple geographic regions and hosts, using `MAFFT` (version 7.490) with default parameters. Phylogenetic trees were generated using the maximum likelihood (ML) method in `RAxML-NG` (version 1.2.2) ^69^, with the best-fit available model selected based on the Bayesian Information Criterion (BIC) values in `IQ-TREE` (version 2.4.0) ^70^.

For Nanopore sequencing data used for mosquito species determination and blood meal analysis, raw data in Fast5 file format from the sequencer were used for base-calling with `Guppy` (version 5.6.7), with a phred quality score ≥ 6 (Q score ≥ 6). Reads were demultiplexed according to their barcodes, which were then trimmed using `Guppy` with defaults. BLAST was applied to all passed reads to investigate the mosquito species and blood sources (calls were made for hits with e-value < 1e-6 and length > 100bp).

All data was visualized in R software (version 4.2.0; R Core Team, Vienna, Austria) in RStudio (version 2022.02.2) with *ggrepel* (version 0.9.6), *ggplot2* (version 3.5.2), *tidyverse* (version 2.0.0), *eulerr* (version 7.0.2), *viridis* (version 0.6.5), *scico* (version 1.5.0), *ape* (version 5.8-1), *ggtree* (version 3.16.0), *rentrez* (version 1.2.4), *countrycode* (version 1.6.1), *RColorBrewer* (version 1.1-3), and *dplyr* (version 1.1.4) packages. Mapping was done using *sf* (version 1.0.21), *rnaturalearth* (version 1.0.1), and *rnaturalearthdata* (version 1.0.0) packages in R software. Data for mapping was downloaded from the Global Administrative Areas Database (GADM) ^71^ and the Humanitarian Data Exchange (HDX) ^72^.

## Acknowledgements

We thank Daniel Neafsey for sharing mosquito blood meal sequencing protocols and advice. The following reagent was provided by the NIH/NIAID Filariasis Research Reagent Resource Center for distribution through BEI Resources, NIAID, NIH: Uninfected *Aedes aegypti*, Strain Black Eye Liverpool (Frozen), NR-48920.

The authors used an artificial intelligence language model for English language editing during the writing process. However, the manuscript is original to the authors, who take responsibility for its content.

## Author contributions

W.H., J.J.J, and J.B.P. conceptualized the study and designed experiments. T.B. and F.V. collected field mosquito samples. M.M.K. and A.K.T. helped coordinate these collections. W.H. performed laboratory tests. W.H., A.P., Z.R.P.-H., and M.H.C. performed data analyses. W.H. prepared the figures and wrote the first draft. All authors reviewed and approved the final draft.

## Data availability

NovaSeq and MinION sequencing data presented in the study are deposited in the NCBI Sequence Read Archive (BioProject ID: PRJNA1200724 and PRJNA1200731).

## Code availability

Analysis code is publicly available at https://github.com/IDEELResearch/DRCAedesSeq.

---

**Figure S1.**
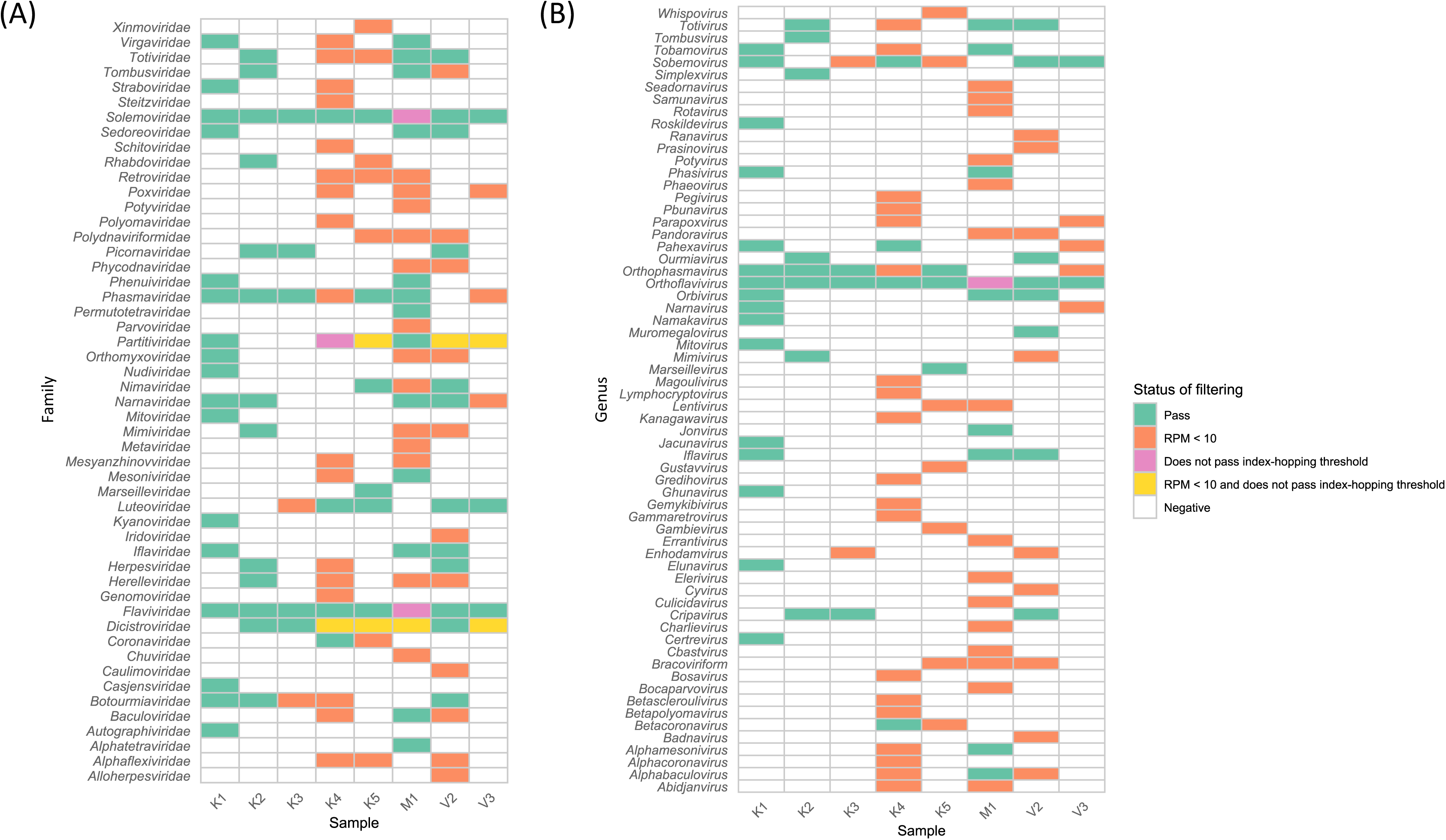
Filtering results by viral family and genus. After applying the RPM < 10 threshold and index-hopping filter, **A)** a total of 33 unique total viral families are detected, with each pool containing between 3-17 passed viral families, and **B)** a total of 28 unique total viral genera are detected, with each pool containing between 2-16 passed viral genera. Shaded cells have detected reads, with color denoting filtering result.

**Figure S2.**
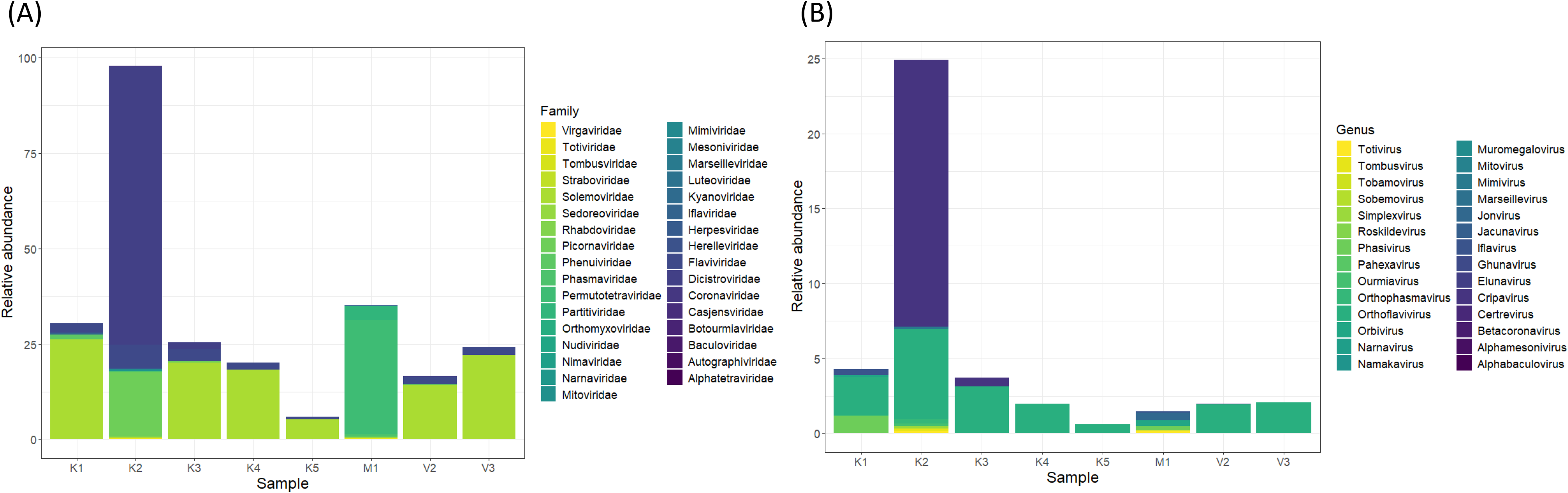
**Viral families and genera detected in field mosquito pools** (<u>after</u> filtering out those with <10 RPM or evidence of index-hopping). **A)** Top five families in each field mosquito pool. **B)** Top five genera in each field mosquito pool. Per International Committee on Taxonomy of Viruses recommendations, relative abundance of *Orthoflavivirus* comprises both *Orthoflavivirus* and *Flavivirus* identified by KrakenUniq.

**Figure S3.**
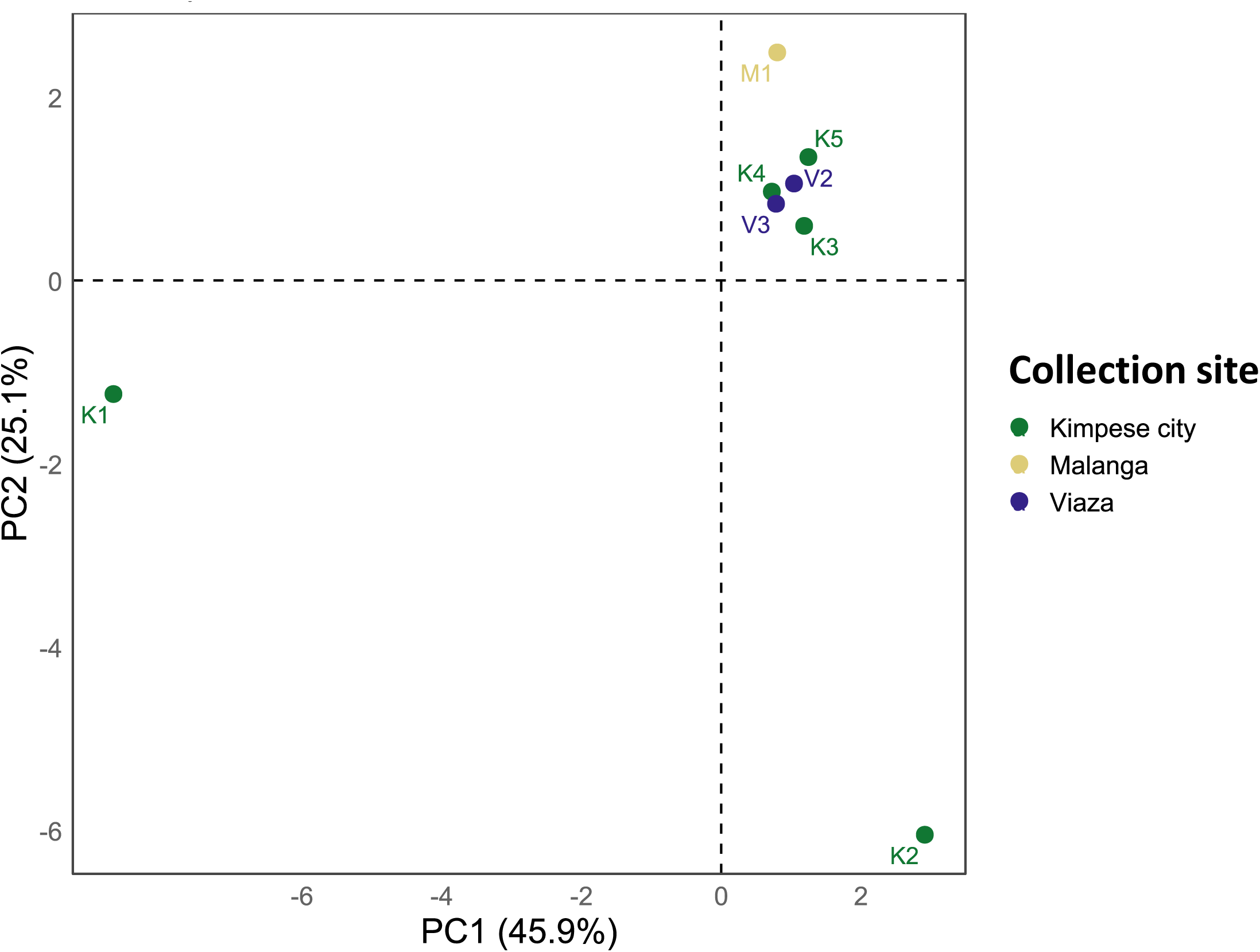
**Principal component analysis (PCA) annotated by collection site** using a data matrix containing relative abundance of viral genera (<u>after</u> applying the 10 RPM threshold and index-hopping filter).

**Figure S4.**
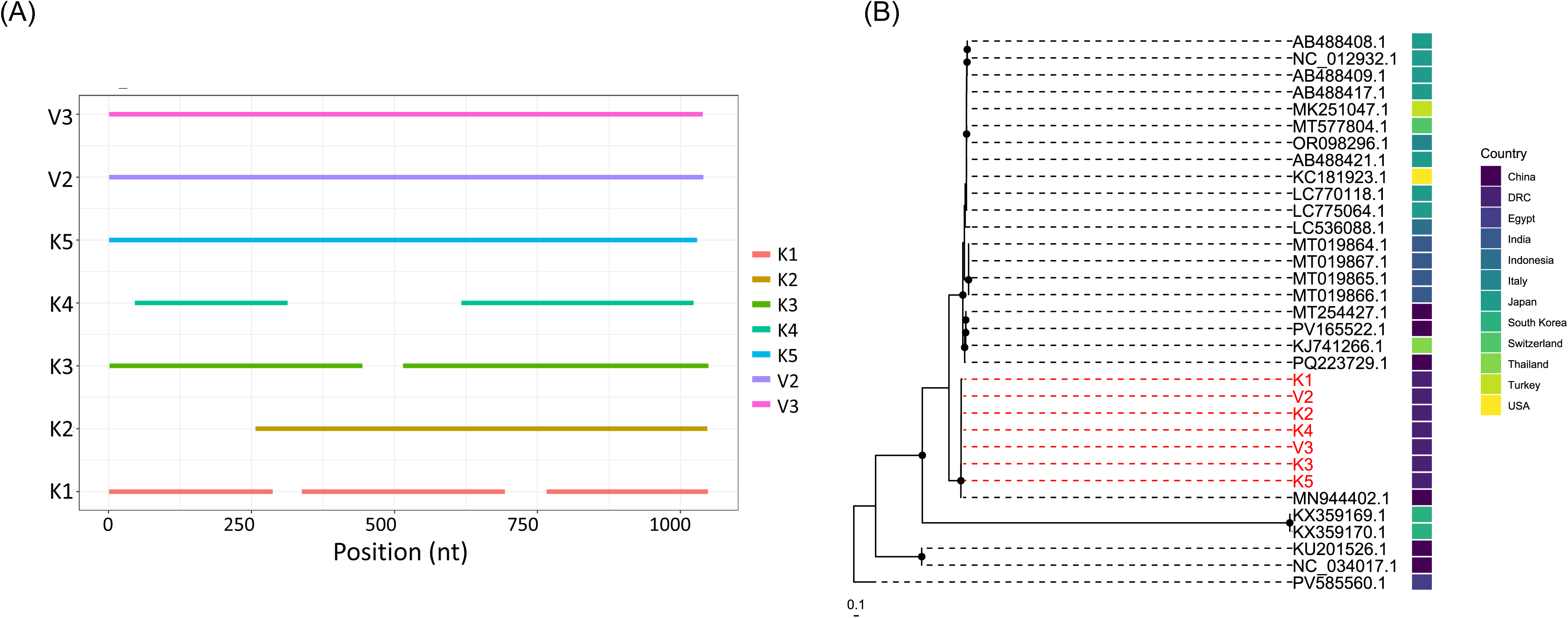
**Coverage plot and phylogenetic analysis of Aedes flavivirus** reads in DRC field mosquito pools. **A)** Coverage of Aedes flavivirus consensus sequences from 7 mosquito pools (x-axis: genomic coordinates of the published Aedes flavivirus sequence MN944402.1), **B)** Aedes flavivirus consensus sequences (red) compared to published sequences (black), constructed using RAxML-NG with the TIM2+F+G4 model and PV585560.1 as outgroup. Analysis of sequences from pools mapping to the polyprotein gene region (MN944402.1, 1-1049, 1049bp). Nodes with >80% bootstrap support across 1,000 replicates are annotated by a black circle.

**Figure S5.**
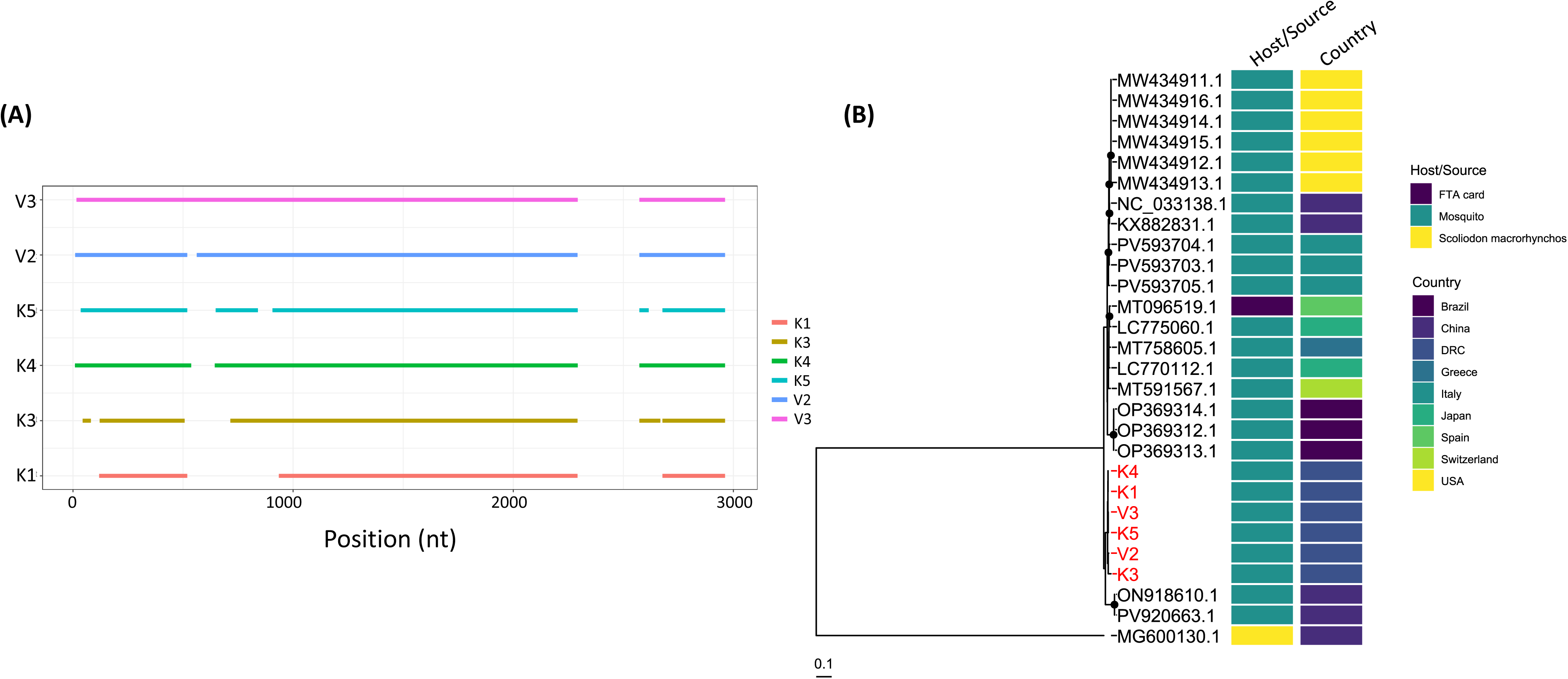
**Coverage plot and phylogenetic analysis of Wenzhou sobemo-like virus 4** reads in DRC field mosquito pools. **A)** Coverage of Wenzhou sobemo-like virus 4 consensus sequences from 6 mosquito pools (x-axis: genomic coordinates of the published sequence ON918610.1), **B)** Wenzhou sobemo-like virus 4 consensus sequences (red) compared to published sequences (black), constructed using RAxML-NG with the SYM+R3 model and MG600130.1 as outgroup. Analysis of sequences from pools mapping to the hypothetical protein 1 (ON918610.1, 69-1380 nt) and hypothetical protein 2 gene (ON918610.1, 1580-2905 nt) regions. Nodes with >80% bootstrap support across 1,000 replicates are annotated by a black circle.

**Figure S6.**
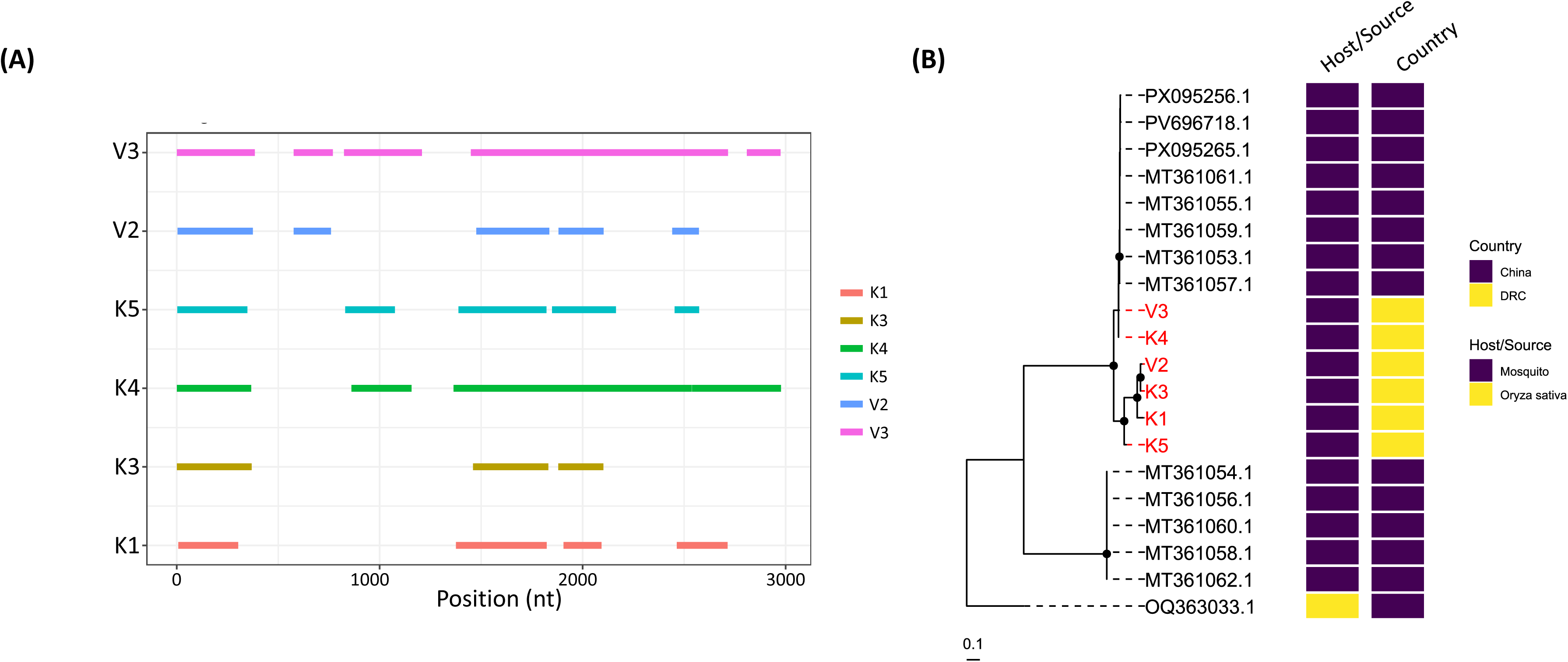
**Coverage plot and phylogenetic analysis of Guangzhou sobemo-like virus** reads in DRC field mosquito pools. **A)** Coverage of Guangzhou sobemo-like virus consensus sequences from 6 mosquito pools (x-axis: genomic coordinates of the published sequence MT361055.1), **B)** Guangzhou sobemo-like virus consensus sequences (red) compared to published sequences (black), constructed using RAxML-NG with the HKY+F model and OQ363033.1 as outgroup. Analysis of sequences from pools mapping to the hypothetical protein (orf1) (MT361055.1, 71-1840 nt) and RNA-dependent RNA polymerase (orf2) gene (MT361055.1, 1582-2907 nt) regions. Nodes with >80% bootstrap support across 1,000 replicates are annotated by a black circle.

**Figure S7.**
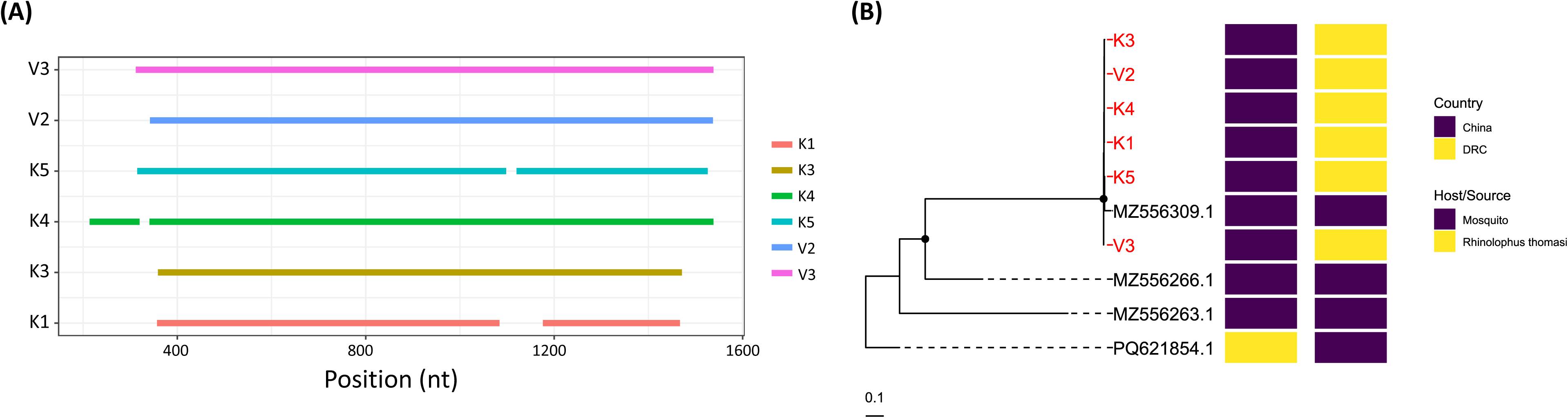
**Coverage plot and phylogenetic analysis of Sichuan mosquito sobemo-like virus** reads in DRC field mosquito pools. **A)** Coverage of Sichuan mosquito sobemo-like virus consensus sequences from 6 mosquito pools (x-axis: genomic coordinates of the published sequence MZ556309.1), **B)** Sichuan mosquito sobemo-like virus consensus sequences (red) compared to published sequences (black), constructed using RAxML-NG with the TPM2+F+I model and PQ621854.1 as outgroup. Analysis of sequences from pools mapping to the capsid protein (MZ556309.1, 214-720 nt) and hypothetical protein gene (MZ556309.1, 823-1538 nt) regions. Nodes with >80% bootstrap support across 1,000 replicates are annotated by a black circle.

**Figure S8.**
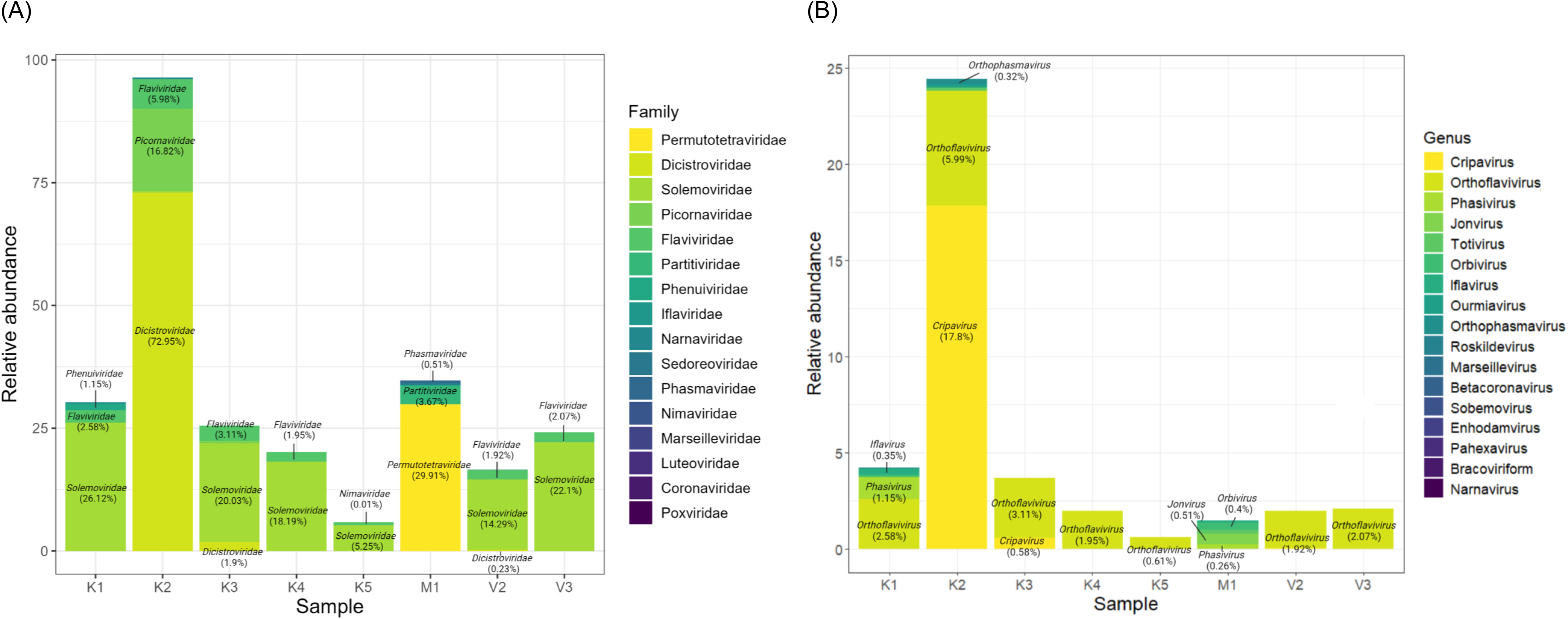
**Viral families and genera in field mosquito pools** (<u>before</u> filtering out those with <10 RPM or evidence of index-hopping). **A)** Top five families in each field mosquito pool; the top three families with relative abundance >0.1% in each pool are labeled. **B)** Top five genera in field mosquito pools; the top three genus with relative abundance >0.1% in each pool are labeled. Per International Committee on Taxonomy of Viruses recommendations, relative abundance of *Orthoflavivirus* comprises both *Orthoflavivirus* and *Flavivirus* identified by KrakenUniq.

**Figure S9.**
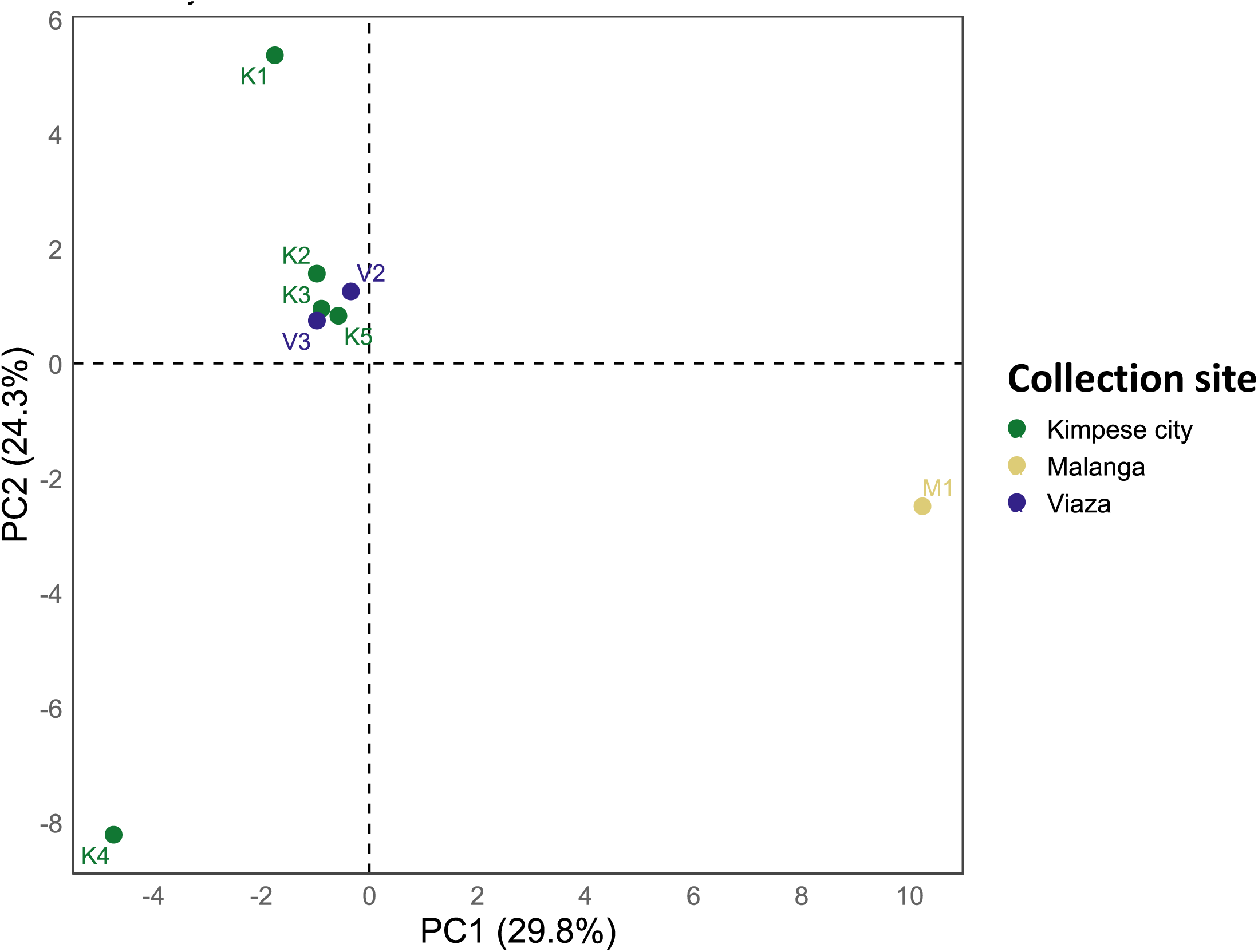
**Principal component analysis (PCA) annotated by collection site** using a data matrix containing relative abundance of viral genera (<u>before</u> applying the 10 RPM threshold and index-hopping filter).

**Figure S10.**
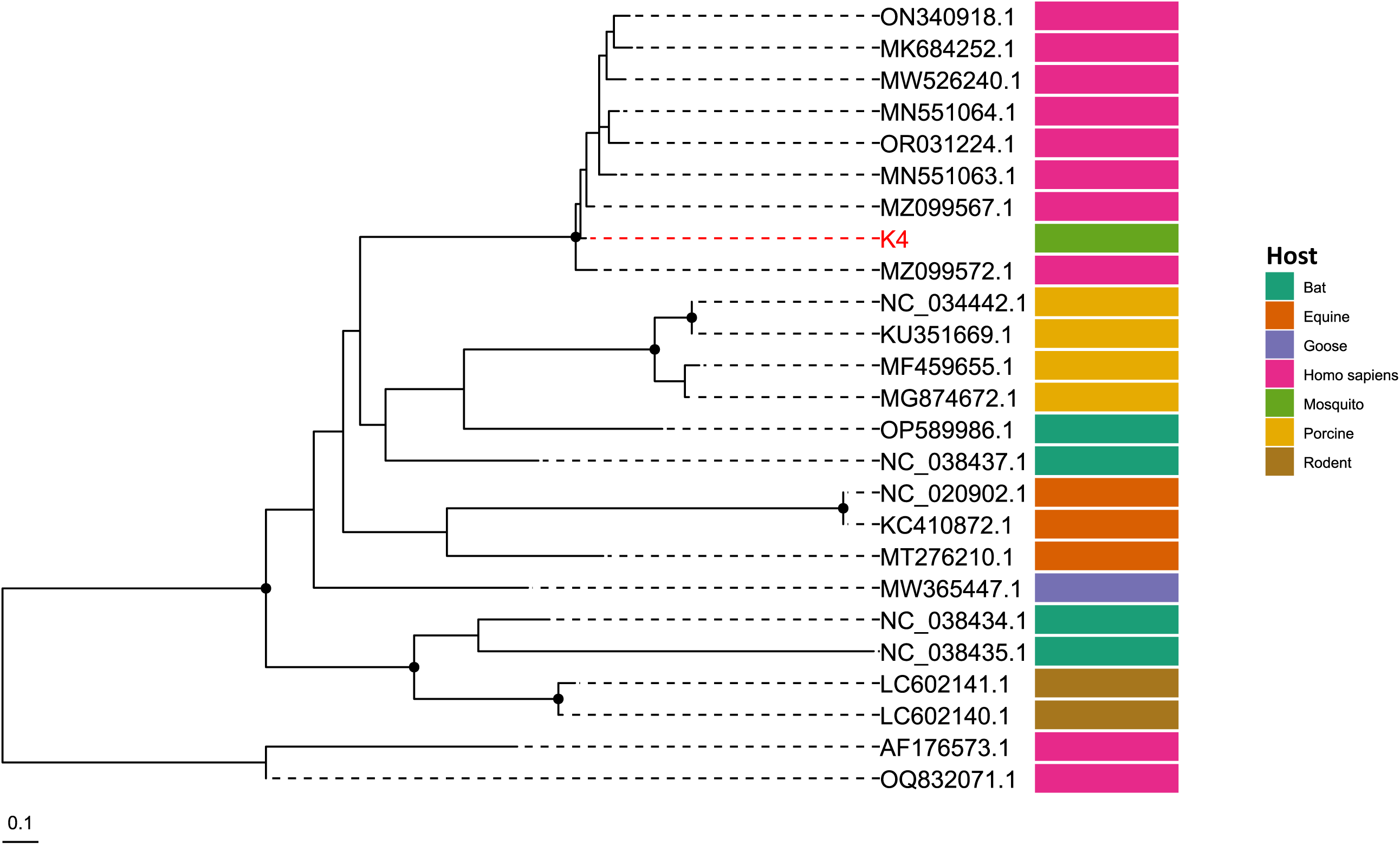
**Phylogenetic analysis of pegivirus sequence** from a DRC field mosquito pool (red) compared to the polyprotein gene region of published sequences (black), constructed using RAxML-NG with the HKY+F+G4 model and hepatitis C virus sequences (AF176573.1 and OQ832071.1) as outgroups. Nodes with >80% bootstrap support across 1,000 replicates are annotated by a black circle.

**Figure S11.**
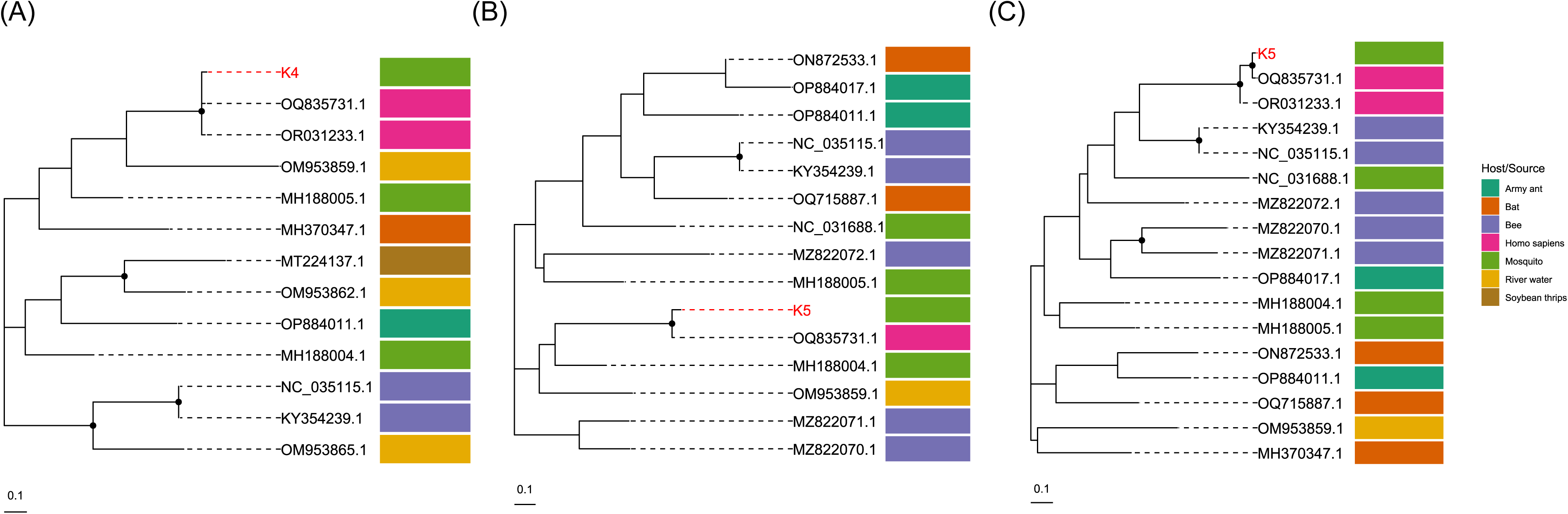
**Phylogenetic analysis of human blood-associated dicistrovirus** sequences in DRC field mosquito pools (red) compared to published sequences (black), constructed using RAxML-NG with the HKY+F+I, HKY+F, and TPM3+F+R2 models, respectively. Analysis of reads from pools **A)** K4 and **B)** K5 mapping to the open reading frame 2 (ORF2) region, and **C)** from K5 mapping to the open reading frame 1 (ORF1) region. Nodes with >80% bootstrap support across 1,000 replicates are annotated by a black circle.

## References

1. Vector-borne diseases. https://www.who.int/news-room/fact-sheets/detail/vector-borne-diseases.

2. Kading, R. C., Brault, A. C. & Beckham, J. D. Global Perspectives on Arbovirus Outbreaks: A 2020 Snapshot. Trop Med Infect Dis 5, (2020).

3. Mbanzulu, K. M. et al. Mosquito-borne viral diseases in the Democratic Republic of the Congo: a review. Parasit. Vectors 13, 103 (2020).

4. Burt, F. J. et al. Chikungunya virus: an update on the biology and pathogenesis of this emerging pathogen. Lancet Infect. Dis. 17, e107–e117 (2017).

5. Tajudeen, Y. A., Oladunjoye, I. O., Mustapha, M. O., Mustapha, S. T. & Ajide-Bamigboye, N. T. Tackling the global health threat of arboviruses: An appraisal of the three holistic approaches to health. Health Promot Perspect 11, 371–381 (2021).

6. Chala, B. & Hamde, F. Emerging and re-emerging vector-borne infectious diseases and the challenges for control: A review. Front. Public Health 9, 715759 (2021).

7. Bobanga, T., Moyo, M., Vulu, F. & Irish, S. R. First Report of *Aedes albopictus* (Diptera: Culicidae) in the Democratic Republic of Congo. Afr. Entomol. 26, 234–236 (2018).

8. Poungou, N. et al. Mosquito-Borne Arboviruses Occurrence and Distribution in the Last Three Decades in Central Africa: A Systematic Literature Review. Microorganisms 12, (2023).

9. Janjoter, S., Kataria, D., Yadav, M., Dahiya, N. & Sehrawat, N. Transovarial transmission of mosquito-borne viruses: a systematic review. Front. Cell. Infect. Microbiol. 13, 1304938 (2023).

10. Renard, A. et al. Interaction of Human Behavioral Factors Shapes the Transmission of Arboviruses by Aedes and Culex Mosquitoes. Pathogens 12, (2023).

11. Kraemer, M. U. et al. The global distribution of the arbovirus vectors and. Elife 4, (2015).

12. Shikanda, H. WHO prequalifies new dengue vaccine, but Africa still not in the picture. Nation https://nation.africa/kenya/health/who-prequalifies-new-dengue-vaccine-but-africa-still-not-in-the-picture-4626442 (2024).

13. Foley, D. H., Rueda, L. M. & Wilkerson, R. C. Insight into global mosquito biogeography from country species records. J. Med. Entomol. 44, 554–567 (2007).

14. Manzambi, E. Z. et al. Behavior of Adult Aedes aegypti and Aedes albopictus in Kinshasa, DRC, and the Implications for Control. Trop. Med. Infect. Dis. 8, (2023).

15. Vulu, F. et al. Geographic expansion of the introduced Aedes albopictus and other native Aedes species in the Democratic Republic of the Congo. Parasit. Vectors 17, 35 (2024).

16. Otshudiema, J. O. et al. Yellow fever outbreak - Kongo Central Province, Democratic Republic of the Congo, August 2016. MMWR Morb. Mortal. Wkly. Rep. 66, 335–338 (2017).

17. Willcox, A. C. et al. Seroepidemiology of Dengue, Zika, and Yellow Fever Viruses among Children in the Democratic Republic of the Congo. Am. J. Trop. Med. Hyg. 99, 756–763 (2018).

18. Selhorst, P. et al. Molecular characterization of chikungunya virus during the 2019 outbreak in the Democratic Republic of the Congo. Emerg. Microbes Infect. 9, 1912–1918 (2020).

19. Mbanzulu, K. M. et al. Mosquito-borne viruses circulating in Kinshasa, Democratic Republic of the Congo. Int. J. Infect. Dis. 57, 32–37 (2017).

20. Sendor, R. et al. Epidemiology of Plasmodium malariae and Plasmodium ovale spp. in Kinshasa Province, Democratic Republic of Congo. Nat. Commun. 14, 6618 (2023).

21. Messina, J. P. et al. The current and future global distribution and population at risk of dengue. Nat Microbiol 4, 1508–1515 (2019).

22. Kajeguka, D. C. et al. Prevalence of dengue and chikungunya virus infections in north-eastern Tanzania: a cross sectional study among participants presenting with malaria-like symptoms. BMC Infect. Dis. 16, 183 (2016).

23. Bernadus, J. B. B. et al. Metagenomic Insight into the Microbiome and Virome Associated with Aedes aegypti Mosquitoes in Manado (North Sulawesi, Indonesia). Infect. Dis. Rep. 15, 549–563 (2023).

24. Gangopadhayya, A. et al. Metagenomic Analysis of Viromes of Aedes Mosquitoes across India. Viruses 16, (2024).

25. Pan, Y.-F. et al. Metagenomic analysis of individual mosquito viromes reveals the geographical patterns and drivers of viral diversity. Nat Ecol Evol 8, 947–959 (2024).

26. Wang, Y. et al. Metagenomic sequencing reveals viral diversity of mosquitoes from Shandong Province, China. Microbiol Spectr e0393223 (2024).

27. Postler, T. S. et al. Renaming of the genus Flavivirus to Orthoflavivirus and extension of binomial species names within the family Flaviviridae. Arch. Virol. 168, (2023).

28. Meng, J.-X. et al. Isolation and genetic evolution of dengue virus from the 2019 outbreak in Xishuangbanna, Yunnan Province, China. Vector Borne Zoonotic Dis. 23, 331–340 (2023).

29. Vogels, C. B. F. et al. DengueSeq: a pan-serotype whole genome amplicon sequencing protocol for dengue virus. BMC Genomics 25, 433 (2024).

30. Fraenkel, S. et al. The development of new primer sets for the amplification and sequencing of the envelope gene of all dengue virus serotypes. Microorganisms 12, 1092 (2024).

31. Rahman, A. et al. Unveiling ancestral relations, host–pathogen interactions and comparative viral miRNA insights of dengue virus serotypes. Network Modeling Analysis in Health Informatics and Bioinformatics 10, 27 (2021).

32. Yung, C.-F. et al. Dengue serotype-specific differences in clinical manifestation, laboratory parameters and risk of severe disease in adults, singapore. Am. J. Trop. Med. Hyg. 92, 999–1005 (2015).

33. Selhorst, P. et al. Phylogeographic analysis of dengue virus serotype 1 and cosmopolitan serotype 2 in Africa. Int. J. Infect. Dis. 133, 46–52 (2023).

34. Garamszegi, L. Z. Host diversity of Aedes albopictus in relation to invasion history: a meta-analysis of blood-feeding studies. Parasit. Vectors 17, 411 (2024).

35. Tchouassi, D. P. et al. Next generation sequencing improves the resolution of detecting mixed host blood meal sources in field collected arboviral mosquito vectors. Med. Vet. Entomol. 38, 407–415 (2024).

36. Almeida, P. R. de, Weber, M. N., Sonne, L. & Spilki, F. R. Aedes-borne orthoflavivirus infections in neotropical primates - Ecology, susceptibility, and pathogenesis. Exp. Biol. Med. 248, 2030–2038 (2023).

37. Hiroshige, Y. et al. A human genotyping trial to estimate the post-feeding time from mosquito blood meals. PLoS One 12, e0179319 (2017).

38. Drummond, C. et al. Stability and detection of nucleic acid from viruses and hosts in controlled mosquito blood feeds. PLoS One 15, e0231061 (2020).

39. Grubaugh, N. D. et al. Xenosurveillance: a novel mosquito-based approach for examining the human-pathogen landscape. PLoS Negl. Trop. Dis. 9, e0003628 (2015).

40. Cordey, S. et al. Detection of dicistroviruses RNA in blood of febrile Tanzanian children. Emerg. Microbes Infect. 8, 613–623 (2019).

41. Oguzie, J. U. et al. Metagenomic surveillance uncovers diverse and novel viral taxa in febrile patients from Nigeria. Nat. Commun. 14, 4693 (2023).

42. Dastjerdi, A., Everest, D. J., Davies, H., Denk, D. & Zell, R. A novel dicistrovirus in a captive red squirrel (Sciurus vulgaris). J. Gen. Virol. 102, (2021).

43. Duraisamy, R. et al. Detection of novel RNA viruses from free-living gorillas, Republic of the Congo: genetic diversity of picobirnaviruses. Virus Genes 54, 256–271 (2018).

44. Asin, I. C. A., Egana, J. M. C., Paul, R. E. & Bautista, M. A. M. Virome sequencing and analysis of Aedes aegypti and Aedes albopictus from ecologically different sites in the Philippines. Parasit. Vectors 18, 426 (2025).

45. Kacnik, S., MacIntyre, C., Guarido, M. & Venter, M. Identification of insect-specific viruses in mosquitoes collected from wildlife and rural areas in north-eastern parts of South Africa using a metagenomic RNA sequencing approach. One Health Outlook 7, 62 (2025).

46. Fumwakwau, J. K., Derra, K., Nzolo, D. B., Mampunza, S. M. M. & Phanzu, D. M. Cohort profile: Kimpese health and demographic surveillance system, democratic republic of Congo. Int. J. Epidemiol. 53, dyae150 (2024).

47. Sun, X. et al. Gut symbiont-derived sphingosine modulates vector competence in Aedes mosquitoes. Nat. Commun. 15, 8221 (2024).

48. Hameed, M. et al. A viral metagenomic analysis reveals rich viral abundance and diversity in mosquitoes from pig farms. Transbound. Emerg. Dis. 67, 328–343 (2019).

49. WHO position paper on dengue vaccines, May 2024. https://www.who.int/publications/i/item/who-wer-9918-203-224.

50. Huang, Y.-M. The Subgenus Stegomyia of Aedes in the Afrotropical Region with Keys to the Species (Diptera: Culicidae). (Magnolia Press, 2004).

51. Phalen, H. et al. A Mosquito Pick-and-Place System for PfSPZ-based Malaria Vaccine Production. IEEE Trans. Autom. Sci. Eng. 18, 299–310 (2020).

52. Zhang, D. X. & Hewitt, G. M. Nuclear integrations: challenges for mitochondrial DNA markers. Trends Ecol. Evol. 11, 247–251 (1996).

53. Batovska, J., Blacket, M. J., Brown, K. & Lynch, S. E. Molecular identification of mosquitoes (Diptera: Culicidae) in southeastern Australia. Ecol. Evol. 6, 3001–3011 (2016).

54. Folmer, O., Black, M., Hoeh, W., Lutz, R. & Vrijenhoek, R. DNA primers for amplification of mitochondrial cytochrome c oxidase subunit I from diverse metazoan invertebrates. Mol. Mar. Biol. Biotechnol. 3, 294–299 (1994).

55. Reeves, L. E. & Burkett-Cadena, N. D. Mosquito blood meal analysis. Cold Spring Harb. Protoc. 2024, db.top107706 (2024).

56. Ogola, E. et al. Composition of Anopheles mosquitoes, their blood-meal hosts, and Plasmodium falciparum infection rates in three islands with disparate bed net coverage in Lake Victoria, Kenya. Malar. J. 16, 360 (2017).

57. Talebzadeh, F. et al. Efficiency of mitochondrial genes and nuclear Alu elements in detecting human DNA in blood meals of Anopheles stephensi mosquitoes: a time-course study. Parasit. Vectors 16, 284 (2023).

58. Meyer, R., Höfelein, C., Lüthy, J. & Candrian, U. Polymerase chain reaction-restriction fragment length polymorphism analysis: a simple method for species identification in food. J. AOAC Int. 78, 1542–1551 (1995).

59. Taylor, P. G. Reproducibility of ancient DNA sequences from extinct Pleistocene fauna. Mol. Biol. Evol. 13, 283–285 (1996).

60. Fauver, J. R. et al. A reverse-transcription/RNase H based protocol for depletion of mosquito ribosomal RNA facilitates viral intrahost evolution analysis, transcriptomics and pathogen discovery. Virology 528, 181–197 (2018).

61. Waggoner, J. J. et al. Single-Reaction Multiplex Reverse Transcription PCR for Detection of Zika, Chikungunya, and Dengue Viruses. Emerg. Infect. Dis. 22, 1295–1297 (2016).

62. Bushnell, B., Rood, J. & Singer, E. BBMerge - Accurate paired shotgun read merging via overlap. PLoS One 12, e0185056 (2017).

63. Bolger, A. M., Lohse, M. & Usadel, B. Trimmomatic: a flexible trimmer for Illumina sequence data. Bioinformatics 30, 2114–2120 (2014).

64. Vasimuddin, M., Misra, S., Li, H. & Aluru, S. Efficient Architecture-Aware Acceleration of BWA-MEM for Multicore Systems. in 2019 *IEEE* International Parallel and Distributed Processing Symposium (IPDPS) 314–324 (IEEE, 2019).

65. Li, H. et al. The Sequence Alignment/Map format and SAMtools. Bioinformatics 25, 2078–2079 (2009).

66. Nurk, S., Meleshko, D., Korobeynikov, A. & Pevzner, P. A. metaSPAdes: a new versatile metagenomic assembler. Genome Res. 27, 824–834 (2017).

67. Breitwieser, F. P., Baker, D. N. & Salzberg, S. L. KrakenUniq: confident and fast metagenomics classification using unique k-mer counts. Genome Biol. 19, 198 (2018).

68. Pan, Y.-F. et al. Metagenomic analysis of individual mosquitos reveals the ecology of insect viruses. bioRxivorg (2023) doi:10.1101/2023.08.28.555221.

69. Kozlov, A. M., Darriba, D., Flouri, T., Morel, B. & Stamatakis, A. RAxML-NG: a fast, scalable and user-friendly tool for maximum likelihood phylogenetic inference. Bioinformatics 35, 4453–4455 (2019).

70. Minh, B. Q. et al. IQ-TREE 2: New models and efficient methods for phylogenetic inference in the genomic era. Mol. Biol. Evol. 37, 1530–1534 (2020).

71. Database of Global Administrative Areas (GADM) (2024). University of California, Berkeley http://www.gadm.org.

72. Welcome - Humanitarian Data Exchange. https://data.humdata.org/.

