## Supplemental Notes for "Evidence of dengue virus transmission and a diverse *Aedes* mosquito virome on the Democratic Republic of Congo-Angola border"

### <u>SUPPLEMENT TO:</u>

**Running title: *Aedes* mosquito virome in the DRC**

Wenqiao He[1#], Thierry Bobanga[2], Anne Piantadosi[3,4], Zachary R. Popkin-Hall[1], Fabien Vulu[2], Matthew H. Collins[4], Melchior M. Kashamuka[5], Antoinette K. Tshefu[5], Jonathan J. Juliano[1*], Jonathan B. Parr[1*#]

[1] Institute for Global Health and Infectious Diseases and Division of Infectious Diseases, Department of Medicine, University of North Carolina at Chapel Hill, Chapel Hill, NC, United States

[2] Department of Tropical Medicine, Faculty of Medicine, University of Kinshasa, Kinshasa, Democratic Republic of the Congo

[3] Department of Pathology and Laboratory Medicine, Emory University School of Medicine, Atlanta, GA, USA

[4] Division of Infectious Diseases, Department of Medicine, Emory University School of Medicine, Atlanta, GA, USA

[5] Kinshasa School of Public Health, Kinshasa, Democratic Republic of the Congo

* Co-senior authors

The main manuscript presents results from metagenomic analysis conducted after applying conservative filters implemented to reduce the risk of false-positive calls. Here, we include additional results of this analysis at the viral family level, as well as results of a separate analysis conducted without the same conservative filters. We have included the latter results because this more lenient approach enables detection of lower frequency viruses within the pools. However, results should be interpreted carefully, with knowledge that there is increased risk of false-positive calls.

**Viral metagenomic analysis at the family level - additional detail (after applying the 10 RPM threshold and an index-hopping filter)**

A total of four viral families—*Iflaviridae*, *Sedoreoviridae*, *Narnaviridae,* and *Totiviridae*—are consistently detected in mosquito pools across all sampling sites. Among these, genera from three families are also present across all sampling sites, including *Iflavirus* (*Iflaviridae*), *Orbivirus* (*Sedoreoviridae*), and *Totivirus* (*Totiviridae*) (**Figure 4**). Reads mapping to *Baculoviridae*, *Alphatetraviridae*, *Mesoniviridae*, and *Permutotetraviridae* are detected in the field mosquito pool collected from Malanga, while none of the pools collected from Kimpese city or Viaza are positive for these viruses. All viral families detected in mosquitoes from Viaza are also present in samples from Kimpese city. *Solemoviridae* exhibit the highest relative abundance in most mosquito pools collected from Kimpese city and Viaza (except for pool K2), while *Permutotetraviridae* is the dominant viral family in mosquitoes from Malanga (**Figure 4 and S2**). The majority of reads within *Solemoviridae* and *Permutotetraviridae* are annotated as insect-specific viruses. Notably, a high relative abundance of *Picornaviridae* and *Dicistroviridae* is observed in one mosquito pool from Kimpese city (K2), with most reads within these families also classified as insect-specific viruses. A relatively high abundance of reads mapping to the *Flaviviridae* family is also observed, except in mosquitoes collected from Malanga and the K2 pool from Kimpese City. Viral families known to include human and animal pathogens and potential pathogens, such as *Flaviviridae, Picornaviridae*, *Rhabdoviridae*, and *Orthomyxoviridae,* are detected.

**Viral metagenomic analysis at the family level - additional detail (prior to applying the 10 RPM threshold and index-hopping filter)**

Based on alignment of the NCBI-nt database, we identify 51 unique viral families in the field mosquito pools, ranging from 7-29 viral families per pool. A total of 17 shared viral families are identified in mosquitoes across all three sample collection areas, with reads mapping to *Solemoviridae* and *Flaviviridae* detected in all field mosquito pools. Similar to the results after applying the RPM <10 threshold and index-hopping filter, the viral family *Solemoviridae* has the highest relative abundance in most of the field mosquito pools collected in Kimpese city and Viaza, while *Permutotetraviridae* has the highest relative abundance in mosquitoes collected in Malanga (**Figure S8a**). Except for the mosquito pool from Malanga, *Flaviviridae* is among the top three viral families in other mosquito pools. High relative abundance of *Picornaviridae* and *Dicistroviridae* is observed in one mosquito pool collected in the Kimpese city (K2).

**Viral metagenomic analysis at the genus level - additional detail (prior to applying the 10 RPM threshold and index-hopping filter)**

We observe a range of 5 to 24 viral genera per field mosquito pool, comprising 64 unique total viral genera. Six viral genera are present in field mosquito pools collected from all sample collection areas, with *Orthoflavivirus* found in all field mosquito pools tested. Similar to the filtered results, *Orthoflavivirus* is the most abundant viral genus in most mosquito pools collected from Kimpese city and Viaza, while the insect virus *Jonvirus* is the genus with the highest relative abundance in the mosquito pool collected in Malanga (**Figure S8b**). PCA based on viral genus annotation to explore clustering by sample collection area is shown in **Figure S9**. Potential geographical clustering is observed among samples from Kimpese city and Viaza. The Malanga mosquito pool is distinct from other samples.

**Phylogenetic analysis of reads mapping to potential pathogens - additional detail (prior to applying the 10 RPM threshold and index-hopping filter)**

BLAST-confirmed reads mapping to viral species related to human or animal diseases, including *human pegivirus* and *human blood-associated dicistrovirus*, are observed in the mosquito pools prior to applying the 10 RPM threshold and index-hopping filter for viral metagenomic data analysis. One consensus sequence showed high similarity with a published human pegivirus polyprotein gene sequence (MZ099567.1, 7917-8114, 228bp) from Thailand was found in a mosquito pool from Kimpese city (**Figure S10**). *Human blood-associated dicistrovirus* reads are detected in two mosquito pools from the Kimpese city. The consensus sequences generated from those reads show high similarity with the structural polyprotein genes (K4 and K5 reads), as well as the non-structural polyprotein (K5 read) gene of an isolate from Cameroon (OQ835731.1, 8846-8978, 133bp;  9276-9430, 155bp;  5259-5406, 148bp) (**Figure S11**).
