## Supplemental Tables for "Evidence of dengue virus transmission and a diverse *Aedes* mosquito virome on the Democratic Republic of Congo-Angola border"

**Table S1.** Mosquito species identified by morphology at each collection site

| Location | *Aedes albopictus* | *Aedes aegypti* | *Aedes simpsoni* | Total |
|---|---|---|---|---|
| Kimpese city | 239 | 1 | 2 | 242 |
| Malanga | 312 | 3 | 1 | 316 |
| Viaza | 102 | 4 | 0 | 106 |
| Total | 653 | 8 | 3 | 664 |

**Table S2.** Mosquito species identification using Nanopore sequencing data

| Sample name | Absolute number of classified reads mapping to *Aedes albopictus* | Absolute number of classified reads mapping to *Aedes aegypti* | Absolute number of classified reads mapping to *Aedes simpsoni* |
|---|---|---|---|
| K1 | 114 | 3 | 0 |
| K2 | 412 | 8 | 0 |
| K3 | 167 | 5 | 0 |
| K4 | 145 | 5 | 0 |
| K5 | 172 | 5 | 0 |
| M1 | 43 | 6 | 0 |
| V2 | 151 | 3 | 0 |
| V3 | 175 | 1 | 0 |

**Table S3.** Microbial genera in the laboratory un-infected *Ae. aegypti* mosquito pool and water control identified using the krakenUniq default nt database (de novo assembled contigs, generated from raw reads after human and mosquito filtering)

| Laboratory mosquito pool | Water control |
|---|---|
| *Chryseobacterium* | *Acinetobacter* |
| *Pseudomonas* | *Cutibacterium* |
| *Enterobacter* | *Herbaspirillum* |
| *Penicillium* | *Methylophilus* |
| *Cupriavidus* | *Corynebacterium* |
| *Aspergillus* | *Gemella* |
| *Azospirillum* | *Hathewaya* |
| *Staphylococcus* | *Malassezia* |
| *Leucobacter* | *Ralstonia* |
| *Wallemia* | *Delftia* |
| *Acinetobacter* | *Inhella* |
| *Variovorax* | *Photorhabdus* |
| *Comamonas* | *Sphingomonas* |
| *Sphingomonas* | *Nocardioides* |
| *Dyadobacter* | *Mycobacterium* |
| *Marinilongibacter* | *Mycolicibacillus* |
| *Siphonobacter* | *Kocuria* |
| *Pedobacter* | *Rothia* |
| *Gordonia* | *Modestobacter* |
| *Rhodococcus* | *Kutzneria* |
| *Betanodavirus* | *Staphylococcus* |
| | *Streptococcus* |
| | *Veillonella* |
| | *Peptoniphilus* |
| | *Cloacibacterium* |
| | *Babesia* |
| | *Natronomonas* |

**Table S4**. Viral genera identified in laboratory un-infected *Ae. aegypti* mosquito pool and water control using KrakenUniq with default nt database (reads could not be *de novo* assembled to contigs)

| Laboratory mosquito pool | Water control |
|---|---|
| *Quaranjavirus* | *Pahexavirus* |
| *Betanodavirus* | *Elvirus* |
| | *Sextaecvirus* |
| | *Andhravirus* |
| | *Schiekvirus* |
| | *Samunavirus* |
| | *Haloferacalesvirus* |
| | *Vojvodinavirus* |
| | *Sanovirus* |
| | *Gammaretrovirus* |
| | *Picobirnavirus* |
| | *Betacoronavirus* |
| | *Allexivirus* |
| | *Parapoxvirus* |
| | *Mimivirus* |
| | *Inovirus* |
| | *Betapapillomavirus* |

**Table S5**. Top ten viral genera contributing to principal components 1 (PC1) and 2 (PC2) during PCA

| PC1 | PC2 |
|---|---|
| *Certrevirus* | *Orthophasmavirus* |
| *Elunavirus* | *Orthoflavivirus* |
| *Ghunavirus* | *Cripavirus* |
| *Jacunavirus* | *Mimivirus* |
| *Mitovirus* | *Simplexvirus* |
| *Namakavirus* | *Tombusvirus* |
| *Narnavirus* | *Ourmiavirus* |
| *Roskildevirus* | *Alphamesonivirus* |
| *Phasivirus* | *Jonvirus* |
| *Iflavirus* | *Alphabaculovirus* |

**Table S6.** DENV read counts, reads per million viral reads identified using the KrakenUniq nt database, and multiplex real-time PCR results using the pan-DENV assay published by Waggoner *et al.* (*Emerg Infect Dis*. 2016)

| Sample | DENV reads, n | DENV reads/million viral reads | Coordinates [reference] | DENV real-time PCR |
|---|---|---|---|---|
| K1 | 1 | 10.3 | 1-90[1] | Negative |
| K2 | 0 | 0 | 0 | Negative |
| K3 | 14 | 31.9 | 1-91[1] | **Positive** |
| K4 | 83 | 68.7 | 1-93[1] | **Positive** |
| K5 | 65 | 27.7 | 1-92[1], 10667-10761[2] | Negative |
| M1 | 0 | 0 | 0 | Negative |
| V2 | 28 | 17.3 | 1-91[1] | Negative |
| V3 | 114 | 32.1 | 1-97[1] | **Positive** |

[1] *Based on published DENV-4 sequence: MG601754.1*
[2] *Based on published DENV-2 sequence: MH048672.1*

**Table S7.** Read counts of *bat faecal associated dicistrovirus 4*

| Sample | *Bat faecal associated dicistrovirus 4* reads, n | Coordinates[1] | Total viral reads (based on nt database) | *Bat faecal associated dicistrovirus 4* reads/million viral reads, n (based on nt database) |
|---|---|---|---|---|
| K1 | 0 | 0 | 96956 | 0 |
| K2 | 33 | 7078-7330 | 2472 | 13349.5 |
| K3 | 182 | 7078-7330 | 439000 | 414.6 |
| K4 | 0 | 0 | 1207424 | 0 |
| K5 | 0 | 0 | 2204216 | 0 |
| M1 | 1 | 0 | 1316904 | 0 |
| V2 | 52 | 7078-7330 | 1618920 | 32.1 |
| V3 | 0 | 0 | 3555364 | 0 |

[1] *Based on published Bat faecal associated dicistrovirus 4 sequence: ON872534.1*

**Table S8.** Read counts of the most prevalent insect-specific viruses (reads/million reads mapping to viruses)

| | K1 | K2 | K3 | K4 | K5 | M1 | V2 | V3 |
|---|---|---|---|---|---|---|---|---|
| *Aedes flavivirus* | 1,320 | 58,252 | 656 | 268 | 230 | 0 | 284 | 71 |
| *Wenzhou sobemo-like virus 4* | 246,875 | 0 | 314,077 | 356,127 | 53,343 | 0 | 211,721 | 320,021 |
| *Hubei mosquito virus 2* | 330 | 0 | 228 | 278 | 76 | 604,697 | 337,739 | 9,509 |
| *Guangzhou sobemo-like virus* | 160,691 | 1,618 | 148,055 | 96,652 | 30,291 | 0 | 103,348 | 145,950 |
| *Sichuan mosquito sobemo-like virus* | 98,890 | 1,618 | 51,417 | 83,487 | 21,927 | 0 | 39,073 | 74,002 |

**Table S9.** Primers for mosquito species determination and blood meal investigation. Reaction mixtures and reaction conditions are the same for all assays.

| Target | Primer sequences 5′ to 3′ | Reaction mixture | Reaction Condition | Study |
|---|---|---|---|---|
| 16S | CGGTTGGGGTGACCTCGGA<br>GCTGTTATCCCTAGGGTAACT | 5 µl 5× PrimeSTAR GXL Buffer (Takara Bio)<br>1 µl PrimeSTAR GXL DNA Polymerase (Takara Bio)<br>4 µl dNTP Mixture (2.5 mM each)<br>200 nM Primer 1<br>200 nM Primer 2<br>2 µl Template<br>Add $H_2O$ to 50 µl | 35 cycles of:<br>98°C for 10 s<br>55°C for 15 s<br>68°C for 1 min;<br>Final extension at 68°C for 2 min;<br>Hold at 4°C | Taylor, 1996 |
| cytB | CCATCCAACATCTCAGCATGATGAAA<br>GCCCCTCAGAATGATATTTGTCCTCA | | | Meyer, Hofelein, Luthy, & Candrian, 1995 |
| COX1 | TACAGTTGGAATAGACGTTGATAC<br>TCCAATGCACTAATCTGCCATATTA | | | Zhang & Hewitt, 1996 |
| COI | GGTCAACAAATCATAAAGATATTGG<br>TAAACTTCAGGGTGACCAAAAAATCA | | | Folmer, 1994 |
| ITS2 | GCTCGTGGATCGATGAAGAC<br>TGCTTAAATTTAGGGGGTGTAGTCAC | | | Batovska, Blacket, Brown, & Lynch, 2016 |

**Table S10**. Sequences for *Bat faecal associated dicistrovirus 4* phylogenetic analysis

| Accession number/ Sample name | Virus name | Host/Source | This study |
|---|---|---|---|
| K2 | Bat faecal associated dicistrovirus 4 | Mosquito | yes |
| K3 | Bat faecal associated dicistrovirus 4 | Mosquito | yes |
| MH188004.1 | Culex dicistrovirus 1 | Mosquito | no |
| MH188005.1 | Culex dicistrovirus 2 | Mosquito | no |
| MT195550.1 | Soybean thrips dicistrovirus 1 | Soybean thrips | no |
| MZ822070.1 | Apis dicistrovirus 2 | Bee | no |
| MZ822071.1 | Apis dicistrovirus 3 | Bee | no |
| OM953862.1 | Flumine dicistrovirus 2 | River water | no |
| OM953865.1 | Flumine dicistrovirus 3 | River water | no |
| ON872534.1 | Bat faecal associated dicistrovirus 4 | Bat | no |
| OQ715884.1 | Wenzhou bat dicistrovirus 3 | Bat | no |
| OQ715887.1 | Wenzhou bat dicistrovirus 6 | Bat | no |
| V2 | Bat faecal associated dicistrovirus 4 | Mosquito | yes |
| MH370347.1 | Bat dicistrovirus | Bat | no |
| OP884017.1 | Army ant associated dicistrovirus 7 | Army ant | no |
| OP884013.1 | Army ant associated dicistrovirus 4 | Army ant | no |
| MT224137.1 | Soybean thrips dicistrovirus 2 | Soybean thrips | no |
| NC_029052.1 | Goose dicistrovirus | Goose | no |

**Table S11**. Sequences for *Aedes flavivirus* phylogenetic analysis

| Accession number/ Sample name | Virus name | Country | This study | Host/Source |
|---|---|---|---|---|
| PV585560.1 | Culex flavivirus | Egypt | no | mosquito |
| PQ223729.1 | Aedes flavivirus | China | no | mosquito |
| OR098296.1 | Aedes flavivirus | Italy | no | mosquito |
| KC181923.1 | Aedes flavivirus | USA | no | mosquito |
| PV165522.1 | Aedes flavivirus | China | no | mosquito |
| LC775064.1 | Aedes flavivirus | Japan | no | mosquito |
| LC770118.1 | Aedes flavivirus | Japan | no | mosquito |
| MT577804.1 | Aedes flavivirus | Switzerland | no | mosquito |
| MT254427.1 | Aedes flavivirus | China | no | mosquito |
| MT019867.1 | Aedes flavivirus | India | no | mosquito |
| MT019866.1 | Aedes flavivirus | India | no | mosquito |
| MT019865.1 | Aedes flavivirus | India | no | mosquito |
| MT019864.1 | Aedes flavivirus | India | no | mosquito |
| LC536088.1 | Aedes flavivirus | Indonesia | no | mosquito |
| NC_012932.1 | Aedes flavivirus | Japan | no | mosquito |
| MK251047.1 | Aedes flavivirus | Turkey | no | mosquito |
| KX359170.1 | Aedes flavivirus | South Korea | no | mosquito |
| KX359169.1 | Aedes flavivirus | South Korea | no | mosquito |
| KJ741266.1 | Aedes flavivirus | Thailand | no | mosquito |
| AB488421.1 | Aedes flavivirus | Japan | no | mosquito |
| AB488417.1 | Aedes flavivirus | Japan | no | mosquito |
| AB488409.1 | Aedes flavivirus | Japan | no | mosquito |
| AB488408.1 | Aedes flavivirus | Japan | no | mosquito |
| NC_034017.1 | Aedes flavivirus | China | no | mosquito |
| KU201526.1 | Aedes flavivirus | China | no | mosquito |
| K1 | Aedes flavivirus | DRC | yes | mosquito |
| K2 | Aedes flavivirus | DRC | yes | mosquito |
| K3 | Aedes flavivirus | DRC | yes | mosquito |
| K4 | Aedes flavivirus | DRC | yes | mosquito |
| K5 | Aedes flavivirus | DRC | yes | mosquito |
| V2 | Aedes flavivirus | DRC | yes | mosquito |
| V3 | Aedes flavivirus | DRC | yes | mosquito |
| MN944402.1 | Aedes flavivirus | China | no | mosquito |

**Table S12**. Sequences for *Wenzhou sobemo-like virus 4* phylogenetic analysis

| Accession number/ Sample name | Virus name | Country | This study | Host/Source |
|---|---|---|---|---|
| MG600130.1 | Wenzhou tombus-like virus 1 | China | no | Scoliodon macrorhynchos |
| OP369314.1 | Wenzhou sobemo-like virus 4 | Brazil | no | Mosquito |
| OP369313.1 | Wenzhou sobemo-like virus 4 | Brazil | no | Mosquito |
| OP369312.1 | Wenzhou sobemo-like virus 4 | Brazil | no | Mosquito |
| PV920663.1 | Wenzhou sobemo-like virus 4 | China | no | Mosquito |
| PV593705.1 | Wenzhou sobemo-like virus 4 | Italy | no | Mosquito |
| PV593704.1 | Wenzhou sobemo-like virus 4 | Italy | no | Mosquito |
| PV593703.1 | Wenzhou sobemo-like virus 4 | Italy | no | Mosquito |
| LC775060.1 | Wenzhou sobemo-like virus 4 | Japan | no | Mosquito |
| LC770112.1 | Wenzhou sobemo-like virus 4 | Japan | no | Mosquito |
| MT591567.1 | Wenzhou sobemo-like virus 4 | Switzerland | no | Mosquito |
| MT758605.1 | Wenzhou sobemo-like virus 4 | Greece | no | Mosquito |
| MW434916.1 | Wenzhou sobemo-like virus 4 | USA | no | Mosquito |
| MW434915.1 | Wenzhou sobemo-like virus 4 | USA | no | Mosquito |
| MW434914.1 | Wenzhou sobemo-like virus 4 | USA | no | Mosquito |
| MW434913.1 | Wenzhou sobemo-like virus 4 | USA | no | Mosquito |
| MW434912.1 | Wenzhou sobemo-like virus 4 | USA | no | Mosquito |
| MW434911.1 | Wenzhou sobemo-like virus 4 | USA | no | Mosquito |
| MT096519.1 | Wenzhou sobemo-like virus 4 | Spain | no | FTA card |
| NC_033138.1 | Wenzhou sobemo-like virus 4 | China | no | Mosquito |
| KX882831.1 | Wenzhou sobemo-like virus 4 | China | no | Mosquito |
| ON918610.1 | Wenzhou sobemo-like virus 4 | China | no | Mosquito |
| K1 | Wenzhou sobemo-like virus 4 | DRC | yes | Mosquito |
| K3 | Wenzhou sobemo-like virus 4 | DRC | yes | Mosquito |
| K4 | Wenzhou sobemo-like virus 4 | DRC | yes | Mosquito |
| K5 | Wenzhou sobemo-like virus 4 | DRC | yes | Mosquito |
| V2 | Wenzhou sobemo-like virus 4 | DRC | yes | Mosquito |
| V3 | Wenzhou sobemo-like virus 4 | DRC | yes | Mosquito |

**Table S13.** Sequences for *Guangzhou sobemo-like virus* phylogenetic analysis

| Accession number/ Sample name | Virus name | Country | This study | Host/Source |
|---|---|---|---|---|
| OQ363033.1 | Hangzhou sobemo-like virus 1 | China | no | Oryza sativa |
| MT361062.1 | Guangzhou sobemo-like virus | China | no | Mosquito |
| MT361061.1 | Guangzhou sobemo-like virus | China | no | Mosquito |
| MT361060.1 | Guangzhou sobemo-like virus | China | no | Mosquito |
| MT361056.1 | Guangzhou sobemo-like virus | China | no | Mosquito |
| MT361055.1 | Guangzhou sobemo-like virus | China | no | Mosquito |
| MT361054.1 | Guangzhou sobemo-like virus | China | no | Mosquito |
| MT361053.1 | Guangzhou sobemo-like virus | China | no | Mosquito |
| PX095265.1 | Guangzhou sobemo-like virus | China | no | Mosquito |
| PX095256.1 | Guangzhou sobemo-like virus | China | no | Mosquito |
| PV696718.1 | Guangzhou sobemo-like virus | China | no | Mosquito |
| MT361059.1 | Guangzhou sobemo-like virus | China | no | Mosquito |
| MT361058.1 | Guangzhou sobemo-like virus | China | no | Mosquito |
| MT361057.1 | Guangzhou sobemo-like virus | China | no | Mosquito |
| K1 | Guangzhou sobemo-like virus | DRC | yes | Mosquito |
| K3 | Guangzhou sobemo-like virus | DRC | yes | Mosquito |
| K4 | Guangzhou sobemo-like virus | DRC | yes | Mosquito |
| K5 | Guangzhou sobemo-like virus | DRC | yes | Mosquito |
| V2 | Guangzhou sobemo-like virus | DRC | yes | Mosquito |
| V3 | Guangzhou sobemo-like virus | DRC | yes | Mosquito |

**Table S14.** Sequences for *Sichuan mosquito sobemo-like virus* phylogenetic analysis

| Accession number/ Sample name | Virus name | Country | This study | Host/Source |
|---|---|---|---|---|
| PQ621854.1 | Yunnan solemo-like virus | China | no | Rhinolophus thomasi |
| MZ556263.1 | Sichuan mosquito sobemo-like virus | China | no | Mosquito |
| MZ556266.1 | Sichuan mosquito sobemo-like virus | China | no | Mosquito |
| K1 | Sichuan mosquito sobemo-like virus | DRC | yes | Mosquito |
| MZ556309.1 | Sichuan mosquito sobemo-like virus | China | no | Mosquito |
| K3 | Sichuan mosquito sobemo-like virus | DRC | yes | Mosquito |
| K4 | Sichuan mosquito sobemo-like virus | DRC | yes | Mosquito |
| K5 | Sichuan mosquito sobemo-like virus | DRC | yes | Mosquito |
| V2 | Sichuan mosquito sobemo-like virus | DRC | yes | Mosquito |
| V3 | Sichuan mosquito sobemo-like virus | DRC | yes | Mosquito |

**Table S15**. Sequences for *Human pegivirus* phylogenetic analysis

| Accession number/ Sample name | Virus name | Host | This study |
|---|---|---|---|
| K4 | Human pegivirus | Mosquito | yes |
| MZ099572.1 | Human pegivirus | Homo sapiens | no |
| ON340918.1 | Pegivirus C | Homo sapiens | no |
| MK684252.1 | Human pegivirus | Homo sapiens | no |
| MW526240.1 | Human pegivirus | Homo sapiens | no |
| MZ099567.1 | Human pegivirus | Homo sapiens | no |
| MN551063.1 | Human pegivirus | Homo sapiens | no |
| OR031224.1 | Pegivirus C | Homo sapiens | no |
| MN551064.1 | Human pegivirus | Homo sapiens | no |
| NC_038437.1 | Pegivirus I | Bat | no |
| MW365447.1 | Goose pegivirus | Goose | no |
| KU351669.1 | Pegivirus suis | Porcine | no |
| NC_034442.1 | Pegivirus K | Porcine | no |
| MF459655.1 | Porcine pegivirus | Porcine | no |
| MG874672.1 | Porcine pegivirus | Porcine | no |
| MT276210.1 | Pegivirus equi | Equine | no |
| NC_020902.1 | Equine pegivirus 1 | Equine | no |
| KC410872.1 | Equine pegivirus 1 | Equine | no |
| NC_038435.1 | Pegivirus G | Bat | no |
| NC_038434.1 | Pegivirus F | Bat | no |
| LC602140.1 | Rodent pegivirus | Rodent | no |
| LC602141.1 | Rodent pegivirus | Rodent | no |
| OP589986.1 | Bat pegivirus | Bat | no |
| AF176573.1 | Hepatitis C virus | Homo sapiens | no |
| OQ832071.1 | Hepatitis C virus | Homo sapiens | no |

**Table S16**. Sequences for *Human blood-associated dicistrovirus* phylogenetic analysis

| Accession number/ Sample name | Virus name | Host/Source | This study |
|---|---|---|---|
| OM953859.1 | Flumine dicistrovirus 1 | River water | no |
| OQ835731.1 | Human blood-associated dicistrovirus | Homo sapiens | no |
| OR031233.1 | Human blood-associated dicistrovirus | Homo sapiens | no |
| MH370347.1 | Bat dicistrovirus | Bat | no |
| NC_035115.1 | Apis dicistrovirus | Bee | no |
| ON872533.1 | Bat faecal associated dicistrovirus 1 | Bat | no |
| OP884011.1 | Army ant associated dicistrovirus 3 | Army ant | no |
| OP884017.1 | Army ant associated dicistrovirus 7 | Army ant | no |
| OQ715887.1 | Wenzhou bat dicistrovirus 6 | Bat | no |
| KY354239.1 | Apis dicistrovirus | Bee | no |
| MH188004.1 | Culex dicistrovirus 1 | Mosquito | no |
| MH188005.1 | Culex dicistrovirus 2 | Mosquito | no |
| MZ822070.1 | Apis dicistrovirus 2 | Bee | no |
| MZ822071.1 | Apis dicistrovirus 3 | Bee | no |
| MZ822072.1 | Apis dicistrovirus 4 | Bee | no |
| NC_031688.1 | Mosquito dicistrovirus | Mosquito | no |
| K4 | Human blood-associated dicistrovirus | Mosquito | yes |
| K5 | Human blood-associated dicistrovirus | Mosquito | yes |
| MT224137.1 | Soybean thrips dicistrovirus 2 | Soybean thrips | no |
| OM953862.1 | Flumine dicistrovirus 2 | River water | no |
| OM953865.1 | Flumine dicistrovirus 3 | River water | no |
